# Structural characterization of an intermediate reveals a unified mechanism for the CLC Cl^−^/H^+^ transport cycle

**DOI:** 10.1101/857136

**Authors:** Tanmay S. Chavan, Ricky C. Cheng, Tao Jiang, Irimpan I. Mathews, Richard A. Stein, Antoine Koehl, Hassane S. Mchaourab, Emad Tajkhorshid, Merritt Maduke

## Abstract

Among coupled exchangers, CLCs uniquely catalyze the exchange of oppositely charged ions (Cl^−^ for H^+^). Transport-cycle models to describe and explain this unusual mechanism have been proposed based on known CLC structures. While the proposed models harmonize many experimental findings, there have remained gaps and inconsistencies in our understanding. One limitation has been that global conformational change – which occurs in all conventional transporter mechanisms – has not been observed in any high-resolution structure. Here, we describe the 2.6 Å structure of a CLC mutant designed to mimic the fully H^+^-loaded transporter. This structure reveals a global conformational change to a state that has improved accessibility for the Cl^−^ substrate from the extracellular side and new conformations for two key glutamate residues. Based on this new structure, together with DEER measurements, MD simulations, and functional studies, we propose a unified model of the CLC transport mechanism that reconciles existing data on all CLC-type proteins.

## INTRODUCTION

CLC transporter proteins are present in intracellular compartments throughout our bodies – in our hearts, brains, kidneys, liver, muscles, and guts – where they catalyze coupled exchange of chloride (Cl^−^) for protons (H^+^) (Jentsch and Pusch, 2018). Their physiological importance is underscored by phenotypes observed in knockout animals, including severe neurodegeneration and osteopetrosis (Sobacchi et al., 1993; Stobrawa et al., 2001; Hoopes et al., 2005; Kasper et al., 2005), and by their links to human disease including X-linked mental retardation, epileptic seizures, Dent’s disease, and osteopetrosis (Lloyd et al., 1996; Hoopes et al., 2005; Veeramah et al., 2013; Hu et al., 2016).

CLC-ec1 is a prokaryotic homolog that has served as a paradigm for the family (Estevez et al., 2003; Lin and Chen, 2003; Engh and Maduke, 2005; Miller, 2006; Matulef and Maduke, 2007). Its physiological function enables resistance to acidic conditions, such as those found in host stomachs (Iyer et al., 2003). Like all CLC proteins, CLC-ec1 is a homodimer in which each subunit contains an independent anion-permeation pathway (Miller and White, 1984; Ludewig et al., 1996; Middleton et al., 1996; Dutzler et al., 2002). Studies of CLC-ec1 revealed the importance of two key glutamate residues – “Glu_ex_” and “Glu_in_” (**Figure 1A**) in the transport mechanism. Glu_ex_ is positioned at the extracellular entryway to the Cl^−^-permeation pathway, where it acts both as a “gate” for the transport of Cl^−^ and as a participant in the transport of H^+^ (Dutzler et al., 2003; Accardi and Miller, 2004). Glu_in_ is located towards the intracellular side of the protein and away from the Cl^−^-permeation pathway, where it appears to act as a proton transfer site (Accardi et al., 2005; Lim and Miller, 2009).

**Figure 1.**
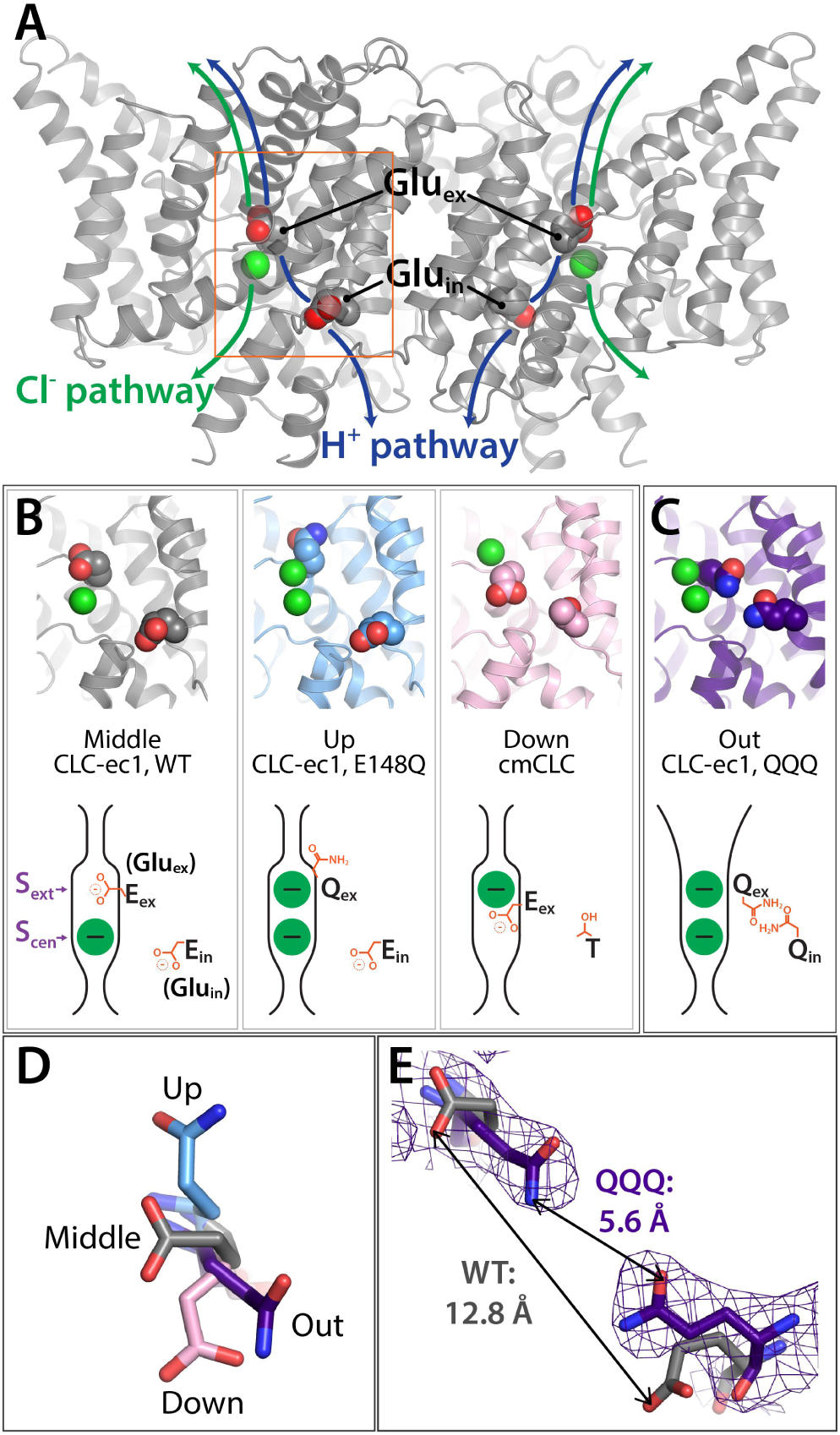
Key glutamate residues in CLC transporters. (**A**) CLC-ec1 wild-type structure (PDB 1ots) showing the external and internal glutamate residues (Glu_ex_ and Glu_in_) in the two subunits of the homodimer. Chloride ions are shown as green spheres. Each subunit independently catalyzes Cl^−^/H^+^ exchange. The approximate transport pathways for these ions are indicated with green and blue arrows. The orange box frames the close-up view (shown in panel B) of the Cl^−^-binding sites along with Glu_ex_ and Glu_in_. (**B**) Three panels showing structures (top panels) and cartoon representations (bottom panels) depicting the three conformations (“middle”, “up”, and “down”) adopted by Glu_ex_ and the position of Glu_in_ (Thr in cmCLC) in various CLC structures, WT (1ots), E148Q (1otu), and cmCLC (3org). The S_ext_ and S_cen_ anion-binding sites are labeled in the WT cartoon at left. (**C**) QQQ structure reveals new conformations for Glu_ex_ and Glu_in_. Structure and cartoon representations as in panel B. (**D**) Overlay of Glu_ex_/Gln_ex_ conformations seen in QQQ (purple), E148Q (blue), WT (grey) and cmCLC (pink). (**E**) Comparison of inter-residue distances and positioning. In WT CLCec1 (grey), the two glutamate residues face away from each other and the distance between them is 12.8 Å. In the QQQ structure, these residues reach inwards to the core of the protein, approaching within 5.6 Å of one another.

Glu_ex_ has been observed in three different positions relative to the Cl^−^-permeation pathway in CLC transporter crystal structures: “middle”, “up”, and “down”. The “middle” conformation is observed in the WT CLC-ec1 structure, where Glu_ex_ occupies the extracellular anion-binding site, “S_ext_” (Dutzler et al., 2002) (**Figure 1B**). The “up” conformation is seen when Glu_ex_ is mutated to Gln, mimicking protonation of Glu_ex_; here, the side chain moves upward and away from the permeation pathway, allowing a Cl^−^ ion to bind at S_ext_ (Dutzler et al., 2003) (**Figure 1B**). The “down” conformation is seen in the eukaryotic cmCLC structure, where Glu_ex_ plunges downwards into the central anion-binding site, “S_cen_” (Feng et al., 2010) (**Figure 1B**). The intracellular anion-binding site, “S_int_”, is a low-affinity site (Picollo et al., 2009) and is not depicted. Of note, cmCLC differs from CLC-ec1 (and from the human CLC transporters) in that it lacks the Glu_in_ residue; a threonine is observed at the corresponding position. In CLC-ec1, a titratable side chain at the Glu_in_ position is required for coupled transport (Lim and Miller, 2009).

The rotation of the Glu_ex_ side chain is the only conformational change that has been detected crystallographically in the CLC transporters. A central question therefore is whether and how other protein conformational changes contribute to the CLC transport mechanism. In previous work, we used a spectroscopic approach to evaluate conformational changes in CLC-ec1, and we found that raising [H^+^] (to protonate Glu_ex_) caused conformational change in regions of the protein outside of the permeation pathway, up to ∼20 Å away from Glu_ex_ (Elvington et al., 2009; Abraham et al., 2015). Using a combination of biochemical crosslinking, double electron-electron resonance (DEER) spectroscopy, functional assays, and molecular dynamics (MD) simulations, we concluded that this H^+^-induced conformational state represents an “outward-facing open” state, an intermediate in the transport cycle that facilitates anion transport to and from the extracellular side (Khantwal et al., 2016).

Here, to obtain a high-resolution structure of the H^+^-induced conformational state, we crystallized a triple mutant, “QQQ”, in which glutamines replace three glutamates: Glu_ex_, Glu_in_, and E113; the latter is a residue within hydrogen bonding distance to Glu_in_ and computationally predicted to be protonated at neutral pH (Faraldo-Gomez and Roux, 2004). In contrast to the single-point mutants of Glu_in_ and Glu_ex_, which reveal either no conformational change (Gln_in_) (Accardi et al., 2005) or only a simple side chain rotation (Gln_ex_) (Dutzler et al., 2003), the QQQ mutant structure reveals global conformational change which generates the expected opening of the extracellular permeation pathway. Unexpectedly, this structure additionally reveals new side chain conformations for both Gln_ex_ and Gln_in_, which bring the two residues to within kissing distance, ∼5 Å. Based on this new structure, together with MD simulations, DEER spectroscopy, and functional studies, we propose an updated model of the Cl^−^ transport cycle.

## RESULTS

### Unique conformations of the H^+^-transfer glutamates

The QQQ mutant (E148Q/E203Q/E113Q) was crystallized in the lipidic cubic phase, without any antibody Fab fragment. The structure, determined at 2.6 Å resolution (**Table 1**), reveals entirely unanticipated changes in the conformations of the Glu_ex_ and Glu_in_ residues, Q148 and Q203. Gln_ex_ residue is in a new position, not previously observed in the CLC transporters, away from the permeation pathway and into the hydrophobic core of the protein, a conformation we designate as “out” (**Figure 1C,D**). Gln_in_, which is positioned identically in all structures to date, is also in a new position, towards the core of the protein. Together, these movements bring Gln_ex_ and Gln_in_ close together, separated by only 5.6 Å (**Figure 1E**). In comparison, this distance is 12.8 Å in the WT CLC-ec1 (1ots) (Dutzler et al., 2003).

**Table 1.**
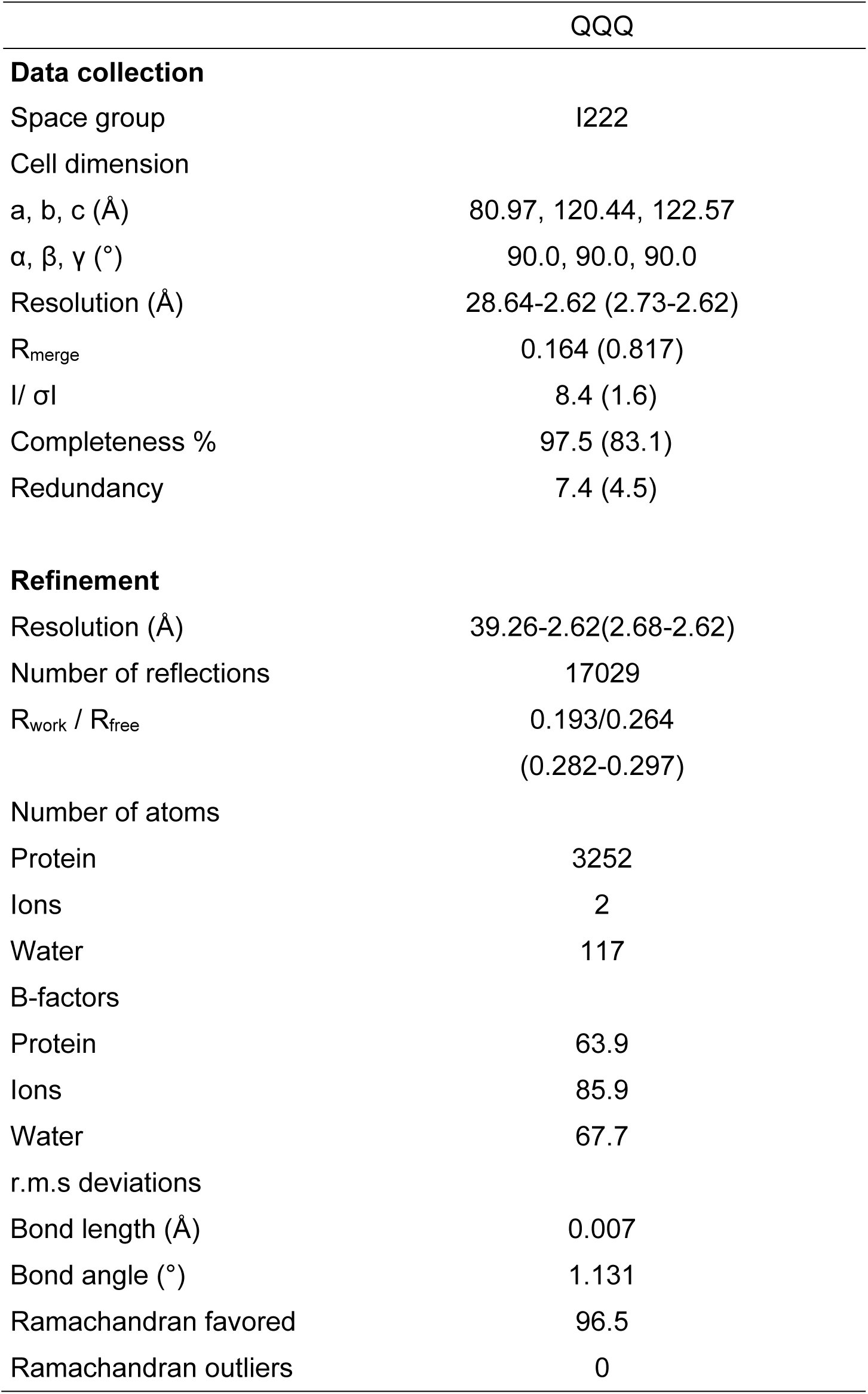
Data collection and refinement statistics. Values in parentheses are for highest-resolution shell

### Opening of the extracellular vestibule and correlation with increased Cl^−^ transport rates

Analysis of the QQQ structure using HOLE, a program for analyzing the dimensions of pathways through molecular structures (Smart et al., 1996), reveals an opening of the extracellular vestibule, increasing accessibility from the extracellular solution to the anion-permeation pathway, in contrast to previously described structures. In the WT protein, two sub-Angstrom bottlenecks occur between S_cen_ and the extracellular side of the protein (**Figure 2A**). In the QQQ protein, these bottlenecks are relieved, widening the pathway to roughly the size of a Cl^−^ ion. In contrast, a single point mutation at the Glu_ex_ position (E148Q) relieves only one of the two bottlenecks (**Figure 2A**). This observation is consistent with the QQQ structure representing the CLC-ec1 outward-facing open state.

**Figure 2.**
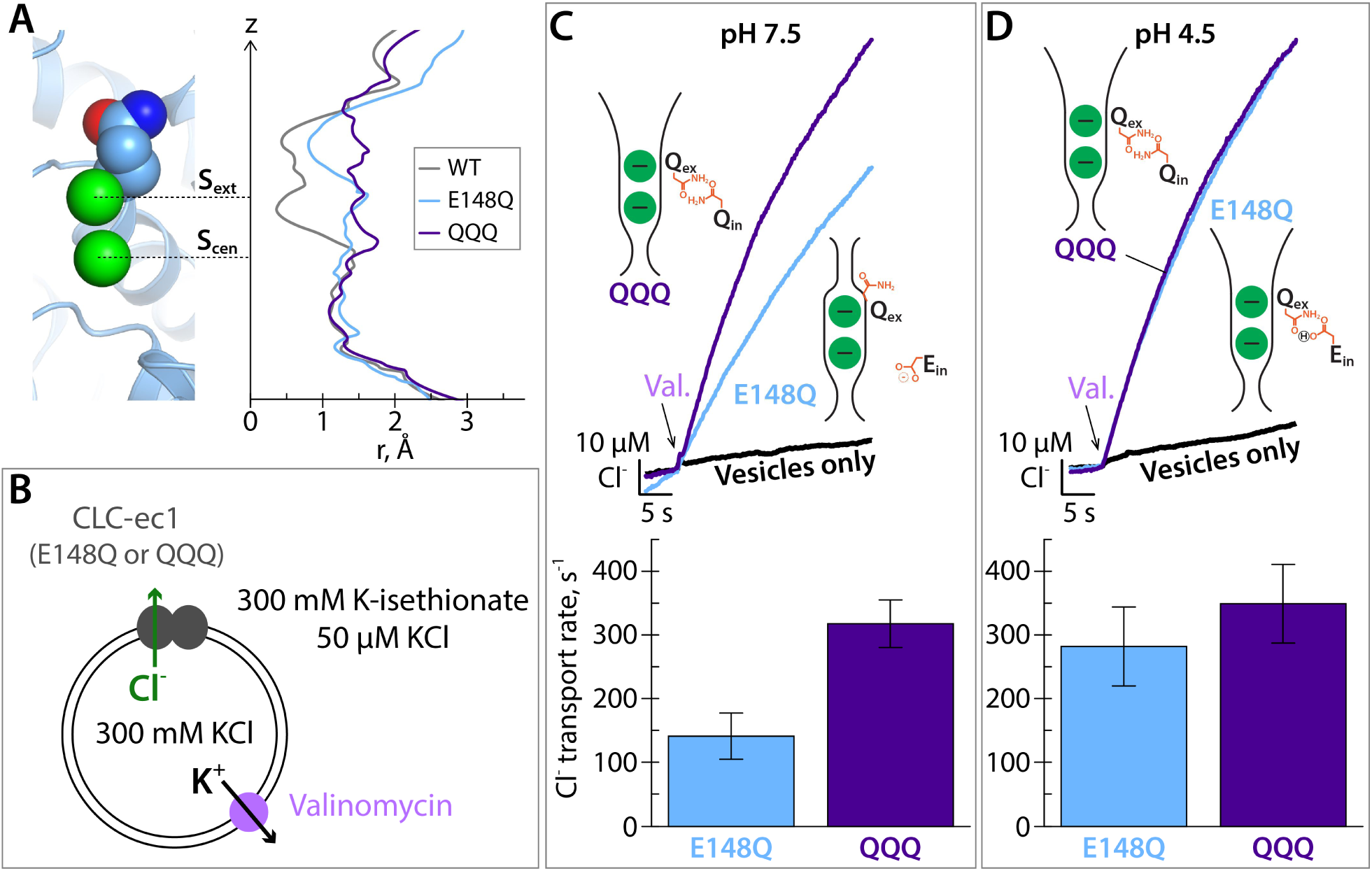
QQQ mutant relieves extracellular bottlenecks for Cl^−^ transport. (**A**) Comparison of pore radius profiles of CLC-ec1 with Glu_ex_ in the “middle” (WT), “up” (E148Q), and “out” (QQQ) positions. The pore-radius profile, calculated using HOLE, is aligned with the structure at left (E148Q) to indicate the position of the S_ext_ and S_cen_ anion-binding sites. (**B**) Cartoon depiction of the Cl^−^ flux assay. The Glu_ex_ mutants E148Q and QQQ catalyze downhill transport of Cl^−^. Bulk movement of Cl^−^ is initiated by the addition of valinomycin, and extravesicular Cl^−^ is quantified using a Ag·AgCl electrode. (**C**) At pH 7.5, Cl^−^ transport through the CLC-ec1 QQQ mutant is ∼2-fold faster compared to E148Q. The top panel shows representative primary data from the flux assay. The bar graph shows summary data from 12 experiments on 3 separate protein preparations. P=0.003 (unpaired t-test). (**D**) At pH 4.5, favoring protonation of glutamate residues, the difference in Cl^−^-transport rates disappears. The bar graph shows summary data from 12 experiments on 3 separate protein preparations. P=0.45 (unpaired t-test).

Mutating Glu_ex_ to Gln renders CLC proteins unable to transport H^+^ but still capable of transporting Cl^−^ (Accardi and Miller, 2004). The difference in the HOLE pore-radius profiles for the QQQ versus E148Q (Glu_ex_ to Gln_ex_) mutants predicts that Cl^−^ transport through QQQ will be faster than through E148Q, assuming no rate-limiting conformational changes intracellular to S_cen_ and S_ext_. Consistent with this prediction, we found Cl^−^ transport through the QQQ mutant to be ∼2-fold faster than through E148Q (**Figure 2B,C**). Since QQQ and E148Q differ at only two residues (glutamine versus glutamate), we hypothesized that lowering the pH to protonate the glutamate residues would allow the E148Q mutant to adopt the QQQ-like (outward-facing open) conformation. In support of this hypothesis, at pH 4.5 there is no difference in transport rates between E148Q and QQQ (**Figure 2D**).

### Structural changes underlying the opening of the extracellular vestibule

The extracellular bottleneck to anion permeation is formed in part by Helix N, which together with Helix F forms the anion-selectivity filter (Dutzler et al., 2002). Previously, we proposed that generation of the outward-facing open state involves movement of Helix N in conjunction with its neighbor Helix P (at the dimer interface) to widen this bottleneck (Khantwal et al., 2016). Structural alignment of the QQQ mutant with either E148Q (**Figure 3A-C**) or WT (**Figure 3C**) confirms the movement of these helices. These structural changes involve shifts in highly conserved residues near the anion-permeation pathway, including F190 (Helix G), F199 (Helix H), and F357 (Helix N). The side chains of all repositioned residues show good electron density (**Figure 3D**, **Figure 3 – figure supplement 1**). Together, these motions widen the extracellular bottleneck (**Figure 3E,F**). This widening is accompanied by subtle changes in the S_ext_ Cl^−^-binding site (**Figure 3 – figure supplement 2A,B**). However, we did not find any significant difference in the affinity of Cl^−^ measured by isothermal titration calorimetry (ITC) (**Figure 3 – figure supplement 2C**), To evaluate changes independent of the structural alignment method, we generated difference distance matrices (Nishikawa, 1972). Comparison of QQQ to WT CLC-ec1 confirms a hot spot of conformational change at Helices M-N-O, as well as changes in Helices C, G, H, and Q (**Figure 3 – figure supplement 3A**). In contrast, comparison of single-Glu mutant structures to WT reveals only minor (≤ 0.8 Å) changes (**Figure 3 – figure supplement 3B**).

**Figure 3.**
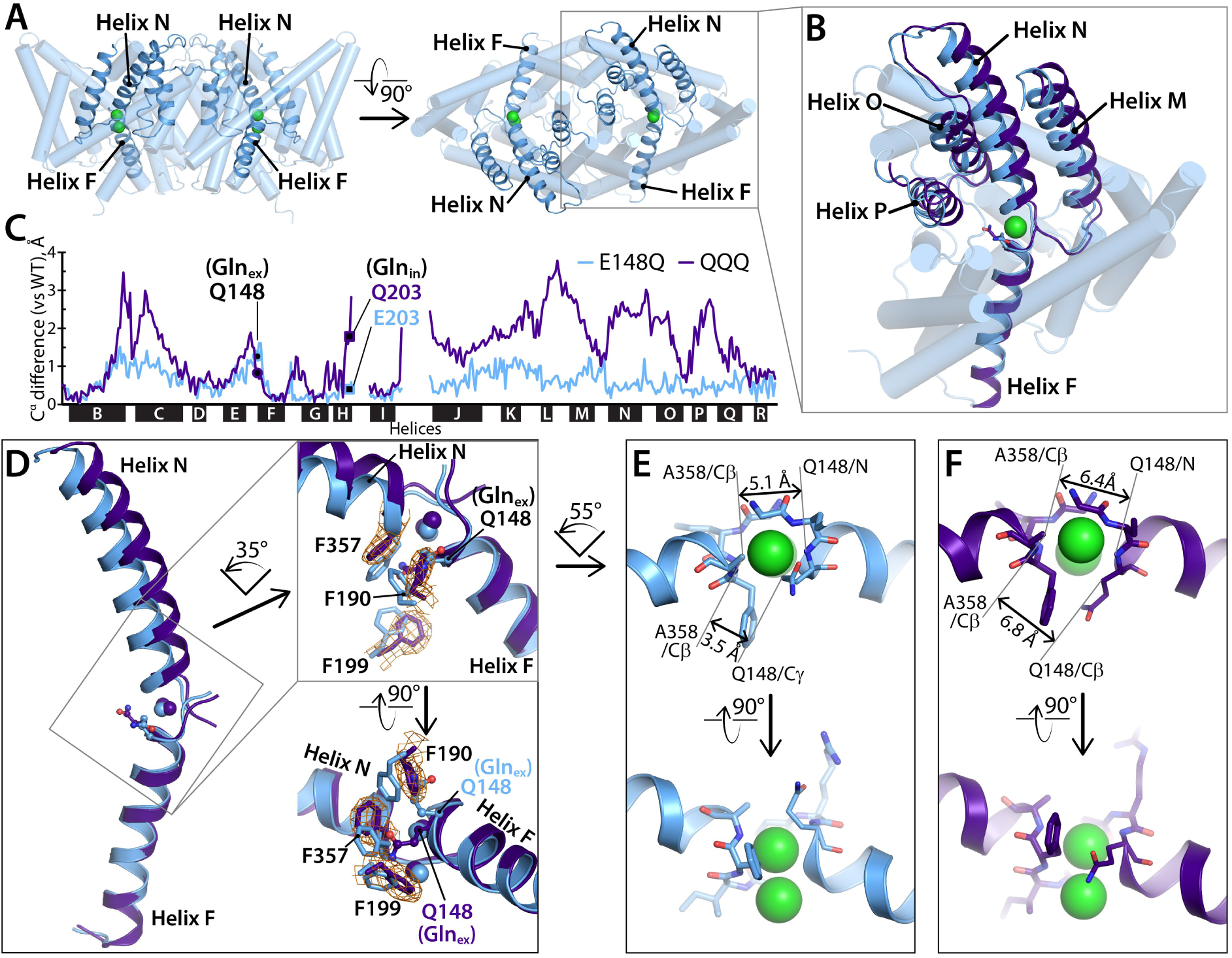
Helices M-N-O-P move to widen the extracellular vestibule. (**A**) Two views of the CLC-ec1 E148Q structure, with ribbons highlighting helices M-N-O-P (helices located in the extracellular half of the protein) and F (located in the intracellular half). (**B**) Overlay of the CLC-ec1 QQQ structure (purple) with E148Q (blue) following structural alignment using segments of helices B, F, G, and I (residues 35-47, 153-164, 174-190, and 215-223). (**C**) Plot of the differences in Cα positions between WT CLC-ec1 and QQQ (purple) or between WT and E148Q (blue), based on the B-F-G-I structural alignment. The black bars indicate 17 of the 18 alpha helices. (Density for Helix A is absent in QQQ.) The locations of Gln_ex_ and Gln_in_ are marked as solid-black circles and squares, respectively. The differences in the H-I and I-J linkers are not shown here but are discussed below and in **Figure 3 – figure supplement 4**. (**D**) Zoomed-in view of the structural overlay between CLC-ec1 QQQ and E148Q, showing changes in the positions of conserved residues F190, F199, and F357 and the corresponding electron density from the QQQ structure determination. Additional views are shown in **Figure 3 – Figure Supplement 1**. (**E, F**) Illustration of interatom distances at the extracellular bottleneck that are increased in QQQ (**F**) compared to E148Q (**E**).

In addition to the conformational changes of the transmembrane helices, there are significant rearrangements of two of the interhelical linkers (**Figure 3 – figure supplement 4A**). At the extracellular side, there are small changes at the I-J linker, which interacts with the Fab fragment in the other structures. At the intracellular side, the Helix H-I linker (residues 205-213) undergoes a several-Angstrom displacement compared to WT. This movement, though relatively large, cannot be of functional significance: previously it was shown that movement of the linker can be restricted via an inter-subunit disulfide cross-link at residue 207, with no significant effect on function (Nguitragool and Miller, 2007). The cross-link at residue 207 does not affect the positioning of Helices H and I (**Figure 3 – figure supplement 4B**).

### Validation of QQQ conformational change in WT CLC-ec1 using DEER spectroscopy

Our working hypothesis is that the QQQ mutant structure mimics the outward-facing open intermediate in the WT CLC transport cycle. When working with a mutant, however, one always wonders whether any conformational change observed is relevant to the WT protein. We therefore used double electron-electron resonance (DEER) spectroscopy to evaluate conformational change in WT CLC-ec1. DEER spectroscopy is advantageous because it can evaluate conformational change by site-directed spin labeling, without the constraints of crystallization. Accurate distance distributions can be obtained for spin labels separated by ∼20-70 Å (Jeschke, 2012; Mishra et al., 2014; Stein et al., 2015). Since CLC-ec1 is a homodimer ∼100 Å in diameter, a simple labeling strategy with one spin label per subunit can provide a sample with optimally spaced probes for distance-change measurements. For example, the extracellular side of Helix N (**Figure 3A**) is separated by ∼50 Å from its correlate in the other subunit. To test the hypothesis that this helix moves, we generated WT CLC-ec1 with spin labels at positions 373 and 374 on Helix N and performed DEER measurements under two conditions, pH 7.5 and pH 4.5. The rationale for this experimental strategy is that pH 4.5 will promote protonation of Glu_ex_ and Glu_in_, thus favoring a global conformation comparable to that observed in the QQQ structure (**Figure 4A**).

**Figure 4.**
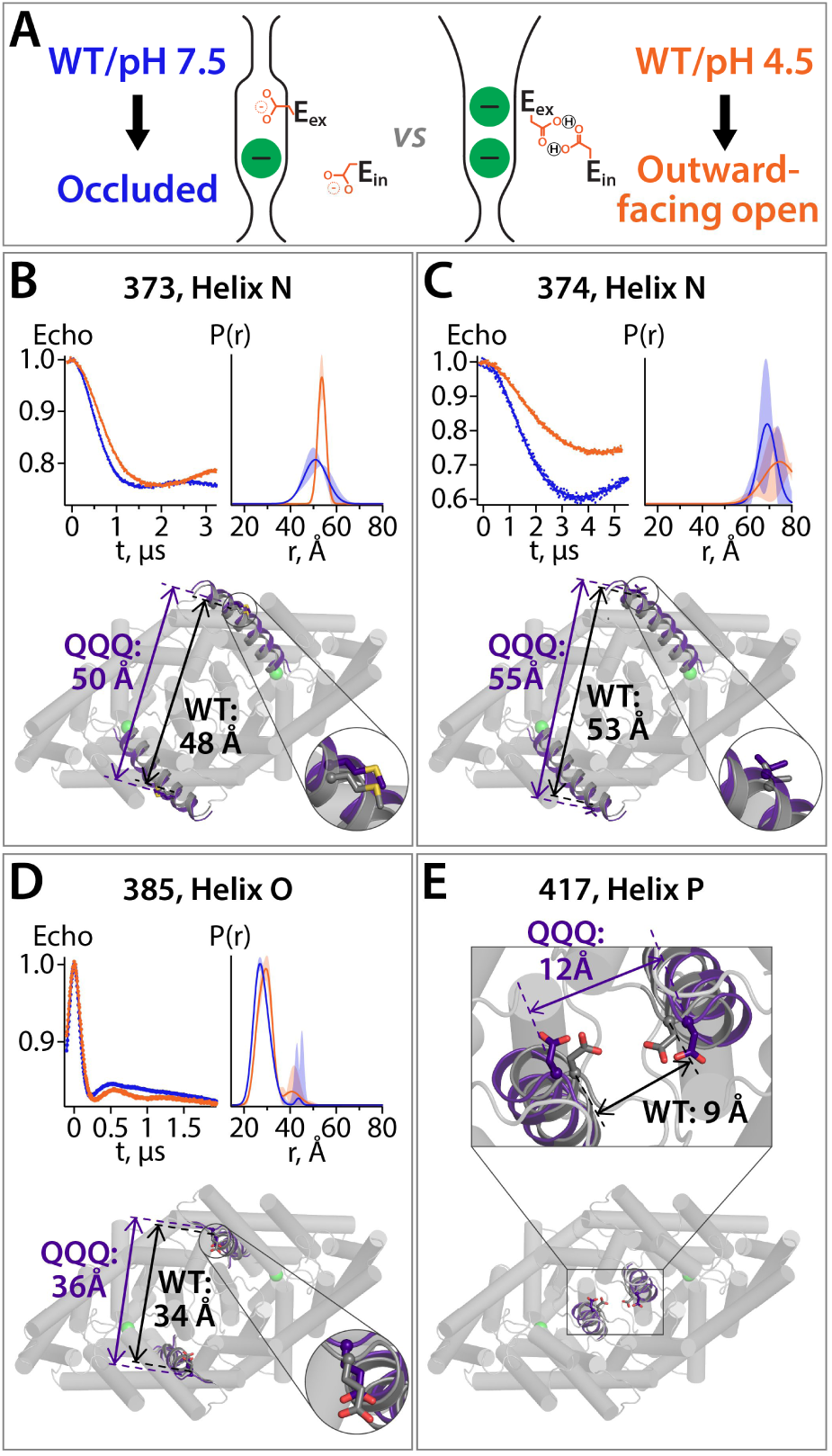
pH-dependent conformational change in WT CLC-ec1. (**A**) Cartoon description of the hypothesis. At pH 7.5, the deprotonated Glu_ex_ and Glu_in_ residues will adopt conformations observed in the WT CLC-ec1 crystal structure, while at pH 4.5 they will adopt conformations observed in the QQQ structure, promoting an overall conformational change that leads to widening of the extracellular vestibule. (**B**) – (**D**) DEER measurements reveal pH-dependent changes in intersubunit distance distributions for spin labels positioned on Helix N or O. The lower panels illustrate the position of each labeled residue on the protein and the Cα intersubunit distances observed in WT versus QQQ. Data for samples with spin labels at residue 373 (B) or 385 (D) were acquired using the standard four-pulse protocol; data for the sample labeled at residue 374 (C) were acquired using the five-pulse protocol (**Figure 4 – figure supplement 2**). (**E**) The intersubunit distance for residue D417 on Helix P increases in QQQ compared to WT. pH-dependent changes in the intersubunit distance for spin-labeled D417C were shown previously (Khantwal et al., 2016).

Consistent with our hypothesis, spin labels on Helix N exhibited pH-dependent changes in distance distributions, in the direction predicted by the QQQ structure (**Figure 4B,C**). Similar results were obtained for a spin label on Helix O (**Figure 4D**). In all cases, spin-labeling did not have any effect on CLC-ec1 activity (**Figure 4 – figure supplement 1**). At Helix P, our previous data provide further support for the relevance of the conformational changes observed in QQQ. First, we showed that CLC-ec1 activity is inhibited by cross-linking of D417C, on Helix P, across the dimer interface (Khantwal et al., 2016). Thus, residues D417 must move apart from one another during the transport cycle. We additionally used DEER to demonstrate that this outward motion of D417C occurs in response to lowering the pH (Khantwal et al., 2016). Correspondingly, the intersubunit Cα distance for D417 is 11.6 Å in the QQQ structure, compared to 8.8 Å in WT (**Figure 4E**), thus supporting the conclusion that the QQQ structure corresponds to a low-pH structure of WT CLC-ec1.

### Analysis of water connections to Gln_ex_

Previous computational studies indicated that water wires can transiently bridge the Glu_ex_ and Glu_in_ residues separated by 12.8 Å in the CLC-ec1 WT structure, which may serve as the pathway for H^+^ transfer (Wang and Voth, 2009; Han et al., 2014). The proximity of these residues in the QQQ structure motivated us to re-evaluate this phenomenon. In our previous studies on WT CLC-ec1, extended MD simulations revealed that water spontaneously enters the hydrophobic core of the protein and transiently and repeatedly forms water wires connecting Glu_ex_ and Glu_in_ (Han et al., 2014). Analogous simulation of the QQQ mutant revealed a dramatic and unanticipated result: water penetration into the hydrophobic core of the protein is greatly increased, and water wires directly connect bulk water in the intracellular solution to Gln_ex_, without requiring intermediate connection to Gln_in_ (**Figure 5A-C**). These water pathways were observed frequently during our 600-ns simulation (**Figure 5D**); in contrast, such water pathways were not observed in our previous 400-ns WT simulation (Han et al., 2014). The number of water molecules needed to reach bulk water follows a normal distribution, with chains of 5 or 6 water molecules predominating (**Figure 5D**). In contrast, the majority of water wires connecting Glu_ex_ to Glu_in_ in the WT simulation involved 7 or more water molecules (Han et al., 2014). Moreover, the occurrence of water pathways in the QQQ simulation (36.5%) is over an order of magnitude greater than the occurrence of water wires between Glu_in_ and Glu_ex_ in the WT simulation (1.3%).

**Figure 5.**
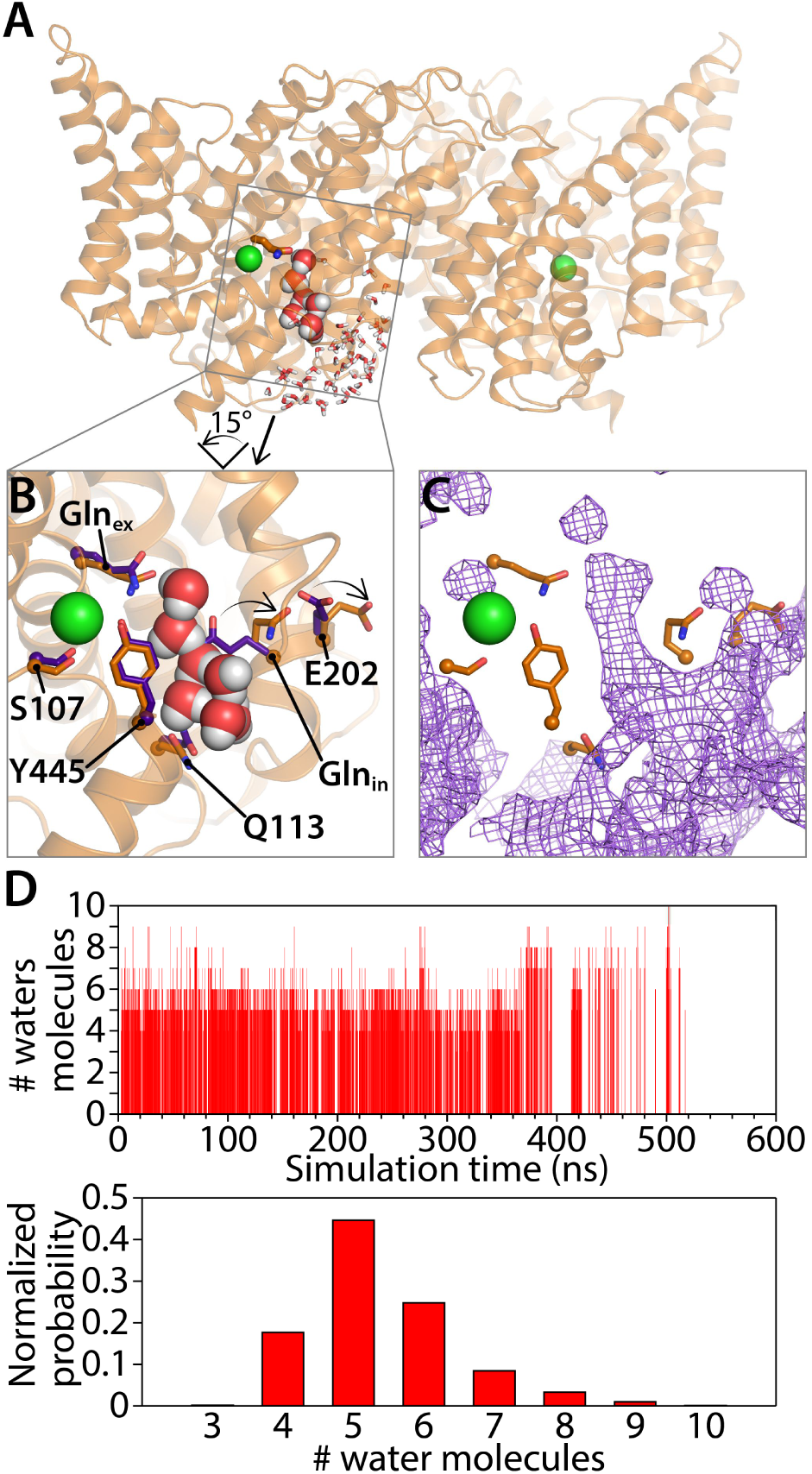
Water pathways between Gln_ex_ and intracellular bulk. (**A**) A simulation snapshot showing a continuous water pathway directly connecting Gln_ex_ to the intracellular bulk water in one of the subunits. (**B**) Zoomed in view of the water pathway. In this example, which represents the most frequently observed water pathway (89% of the water pathways), Gln_in_ has rotated away from the position observed in the QQQ crystal structure to make room for the water pathway. The purple side chains show the residue positions observed in the crystal structure; the orange side chains show the residue positions in a representative simulation snapshot. Conserved residue E202 also rotates from its starting position. (**C**) Overall water occupancy map (density contoured at isovalue 0.35) for the side chain configuration shown in panel B. (**D**) Water pathways between Gln_ex_ and the intracellular bulk water arise spontaneously throughout the 600-ns simulation. Each vertical line in the panels shows the occurrence of a Gln_ex_/bulk-connecting water pathway at that time point, with the length of the line representing the number of water molecules in the shortest path. The number of water molecules needed to reach the bulk water (summarized below) follows a normal distribution dominated by 5-6 water molecules.

The absence of water pathways in the WT simulation is likely due to steric hindrance by Glu_in_, E113, and bulky side chains in the vicinity, which together block direct access of intracellular bulk water toward the protein interior (despite the conformational flexibility of Glu_in_ (Wang and Voth, 2009)). In the QQQ simulation, Gln_in_ can equilibrate among five side chain conformations (Clusters 1-5), all of which can support water pathways (**Figure 5 – figure supplement 1A,B**). Most of the water pathways (96% of pathways observed) occur when Gln_in_ is rotated away from its starting conformation (Clusters 1-3), allowing water to flow along a pathway near Q113 (**Figure 5 – figure supplement 1C,D**). In these conformations, the Gln_in_ side chain bends away from Q113 and from the bulky residues F199 and I109, thus allowing intracellular bulk water to enter the protein interior without encountering steric occlusion (**Figure 5 – figure supplement 2A**).

The predominant water pathway observed in our simulations is roughly parallel to the Cl^−^ permeation pathway (**Figure 5A**). This pathway for water (and hence H^+^) entry into the protein is different from that previously suggested by us and others. Previously, it was proposed that H^+^ access to the interior of the protein occurs via an entry portal located near the interfacial side of the homodimer (Lim et al., 2012; Han et al., 2014) rather than on the “inner” pathway observed here. While we do see some water pathways occurring along the interfacial route, on a pathway that is lined by Gln_in_, these occur only rarely (**Figure 5 – figures supplement 1D**). Importantly, the previous mutagenesis studies supporting the interfacial route are also concordant with the inner water pathway observed here. In the previous studies, mutations that add steric bulk at either E202 (Lim et al., 2012) or the adjacent A404 (Han et al., 2014) were found to inhibit the H^+^ branch of the CLC-ec1 transport cycle. The observation that all water pathways involve rotation of E202 away from its starting position (**Figure 5B**, **Figure 5 – figure supplement 2B, Figure 5 – figure supplement 3**), can explain why bulky mutations at this position would interfere with H^+^ transport.

### Proton pumping without a titratable residue at the Glu_in_ position

Glu_in_ has long been modeled as a H^+^-transfer site in the CLC Cl^−^/H^+^ mechanism (Accardi et al., 2005; Miller, 2006; Lim and Miller, 2009; Basilio et al., 2014; Accardi, 2015; Khantwal et al., 2016). However, this modeling is contradicted by the observation that several CLC transporters can pump H^+^ in the absence of a titratable residue at the Glu_in_ position (Feng et al., 2010; Phillips et al., 2012; Stockbridge et al., 2012). In our analysis of water pathways in the MD simulations, we observed that these pathways are not always lined by the Gln_in_ side chain (**Figure 5 – figure supplement 1**). This finding strongly suggests that while Glu_in_ *facilitates* water pathways, it is not required as a direct H^+^- transfer site. To test this hypothesis experimentally, we evaluated for Cl^−^-coupled H^+^ pumping in two mutants lacking a titratable residue at Glu_in_. In the first experiment, we replaced Glu_in_ (E203) and its H-bonding partner, E113, with the residues found in cmCLC, a eukaryotic transporter that catalyzes 2:1 stoichiometric exchange of Cl^−^ for H^+^ (Feng et al., 2010). In support of our hypothesis, we found that the E203T/E113K mutant catalyzes Cl^−^-driven H+ pumping (**Figure 6A-C**). While the coupling stoichiometry is substantially degraded compared to the WT protein (**Figure 6D**), the thermodynamic fact arising from this experiment is that H^+^ pumping occurs with a non-titratable (Thr) residue at Glu_in_. By comparison, in line with conventional modeling, the Glu_ex_ mutant E148Q wholly fails to catalyze proton pumping (**Figure 6A-C**). To test whether H^+^ pumping requires polar residues at Glu_in_ and/or E113 positions, we examined the double alanine mutant, E203A/E113A. Even with two alanines at these positions, H^+^ pumping is retained (**Figure 6A-C**). Thus, conventional thinking of Glu_in_ as a H^+^- transport site must be re-evaluated.

**Figure 6.**
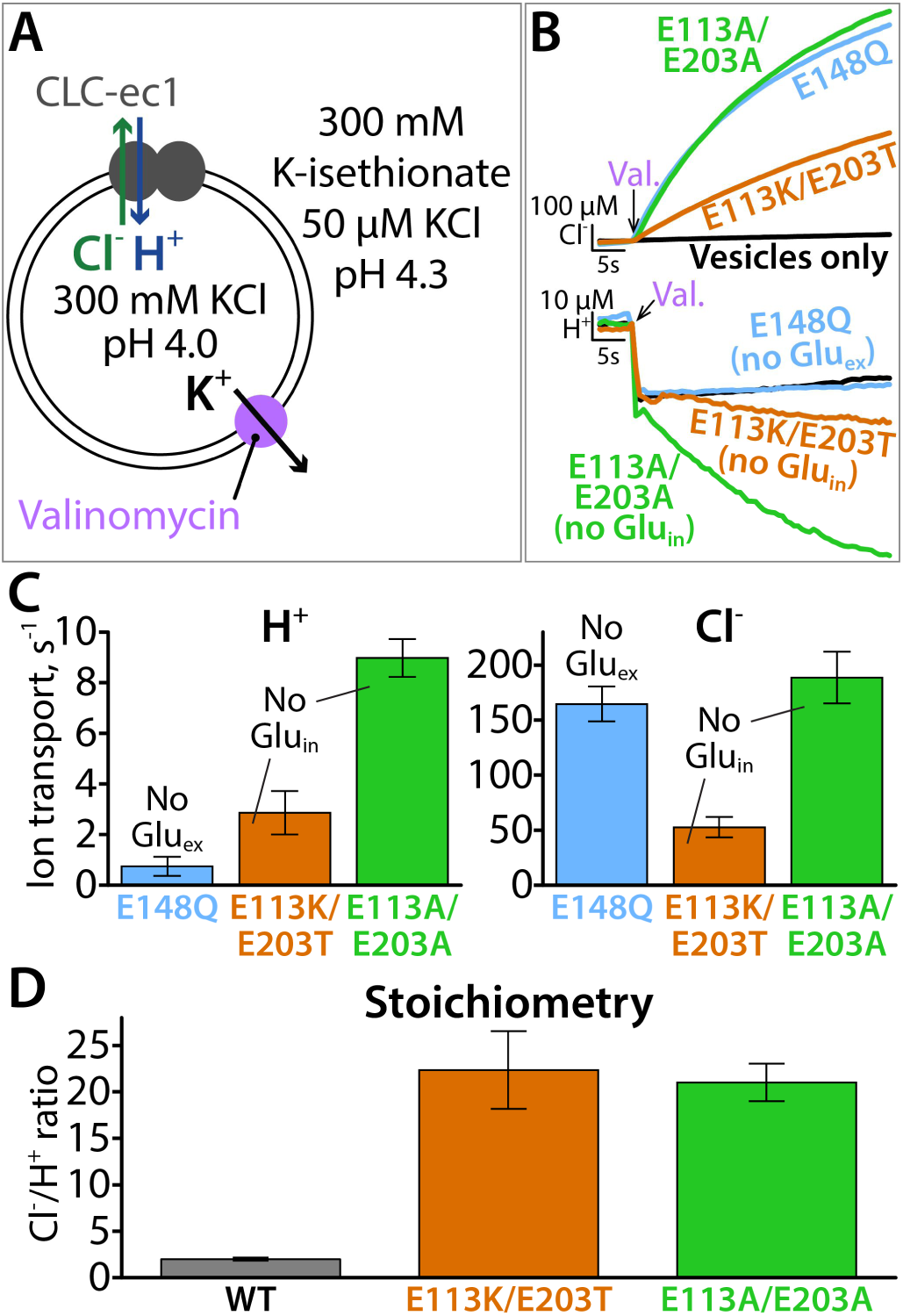
Proton pumping by CLC-ec1 variants with neutral residues at the Glu_in_ position. (**A**) Cartoon depiction of the H^+^/Cl^−^ flux assay. Extravesicular [Cl^−^] and [H^+^] are simultaneously measured using Ag·AgCl and pH electrodes, respectively. The experimental setup involves a 2-fold gradient for H^+^, such that any leak will involve movement of H^+^ out of the vesicles, and any H^+^ movement into the liposomes must occur via transport coupled to the Cl^−^ gradient (pumping). (**B**) Representative Cl^−^ and H^+^ traces for CLC-ec1 mutants with neutral residues at Glu_in_ (residue h203: E113K/E203T and E113A/E203A) or at Glu_ex_ (E148Q). (**C**) Summary data showing H^+^ and Cl^−^ transport rates, average ± SEM. For each mutant, n = 6, with samples from 2 independent protein preparations. (**D**) Cl^−^/H+ stoichiometry for the Glu_in_ mutants, with WT CLC-ec1 shown for comparison. Stoichiometry is determined from the ratio of the ion transport rates (panel C). WT samples were reconstituted at the same time as mutant samples, n = 6 from one protein preparation.

### A unifying model for the CLC transport mechanism

Based on the information gleaned from our study of the QQQ structural intermediate, we propose an updated transport model for 2:1 Cl^−^/H^+^ exchange by CLC transporters. This updated model is inspired by four key findings. First, the outward-facing state has improved accessibility for Cl^−^ to exchange to the extracellular side (**Figure 2**). This state had been previously predicted (Khantwal et al., 2016) but is now seen in molecular detail. Second, the protonated Glu_ex_ can adopt an “out” conformation, within the hydrophobic core of the protein (**Figure 1D**). This novel conformation allows us to eliminate a disconcerting step that was part of all previous models: movement of a protonated (neutral) Glu_ex_ – in competition with Cl^−^ – into the S_cen_ anion-binding site (Miller and Nguitragool, 2009; Feng et al., 2012; Basilio et al., 2014; Khantwal et al., 2016). Third, water pathways can connect Glu_ex_ directly to the intracellular solution (**Figure 5**). Finally, H^+^ pumping does not require a titratable residue at Glu_in_ (**Figure 6**). Together, these findings allow us to propose a revised Cl^−^/H^+^ exchange model that maintains consistency with previous studies and resolves lingering problems.

In our revised model (**Figure 7A**), the first three states are similar to those proposed previously (Miller and Nguitragool, 2009; Feng et al., 2012; Basilio et al., 2014; Khantwal et al., 2016). State A reflects the structure seen in WT CLC-ec1, with Glu_ex_ in the “middle” conformation, occupying S_ext_, and a Cl^−^ occupying S_cen_. Moving clockwise in the transport cycle, binding of Cl^−^ from the intracellular side displaces Glu_ex_ by a “knock-on” mechanism (Miller and Nguitragool, 2009), pushing it to the “up” position and making it available for protonation from the extracellular side (State B). Protonation generates state C, which reflects the structure seen in E148Q CLC-ec1 where Gln_ex_ mimics the protonated Glu_ex_. This sequence of Cl^−^ binding and protonation is consistent with the experimental finding that Cl^−^ and H^+^ can bind simultaneously to the protein (Picollo et al., 2012). Subsequently, a protein conformational change generates an “outward-facing open” state (D). While this state had previously been postulated (Khantwal et al., 2016), the QQQ structure presented here provides critical molecular details.

**Figure 7.**
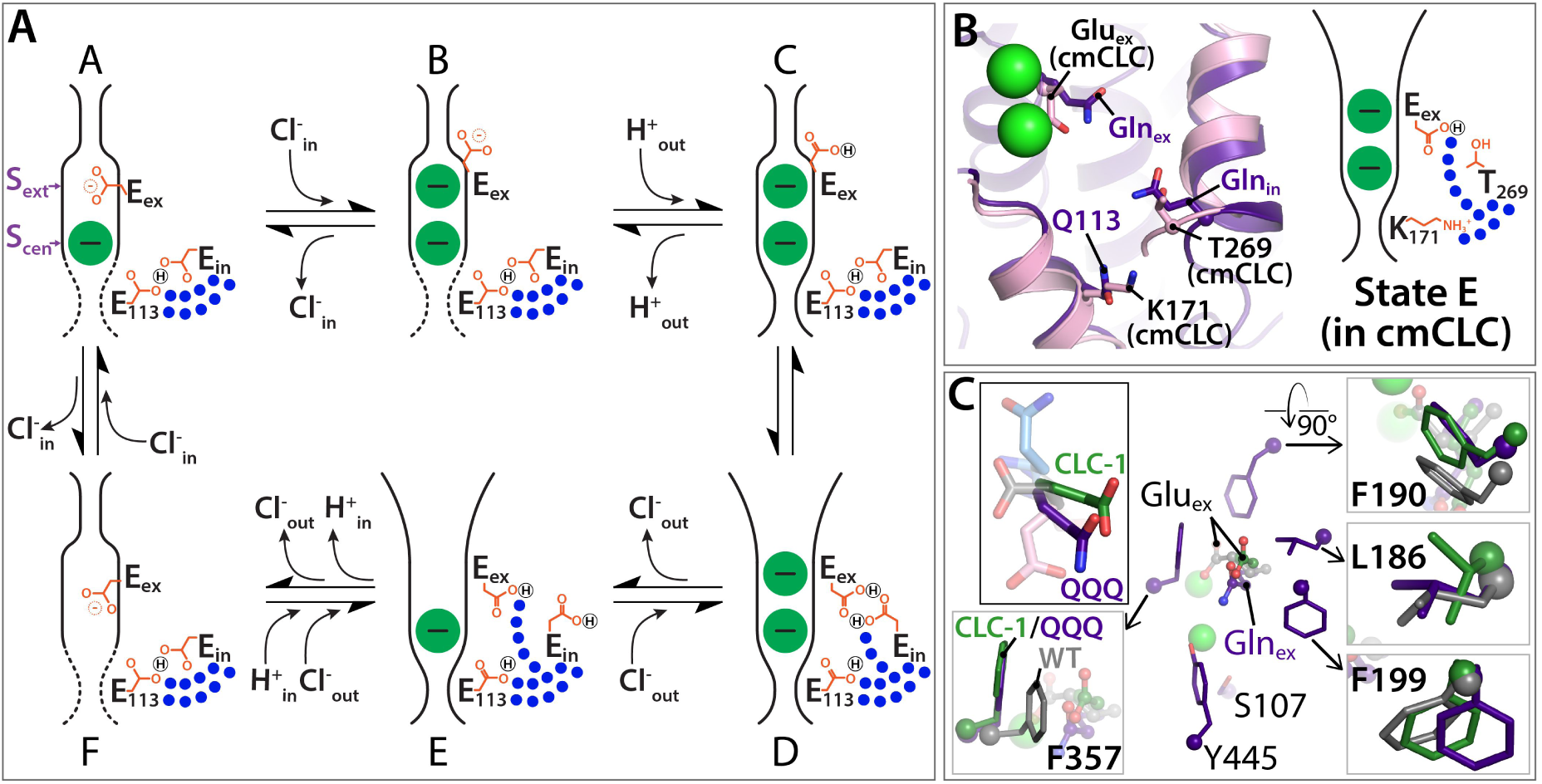
Updated model for the CLC Cl^−^/H^+^ transport cycle (a) Starting from state A, which reflects the structure seen in WT CLC-ec1, a single Cl^−^ is bound, and Glu_ex_ is in the “middle” position. Glu_in_ and E113 are in a H-bonded configuration, restricting water access to the center of the protein. Moving clockwise, binding of Cl^−^ from the intracellular side displaces Glu_ex_ by a “knock-on” mechanism (Miller and Nguitragool, 2009), making it available for protonation from the extracellular side (State B). This protonation step generates state C, which reflects the structure seen in E148Q CLC-ec1, with Glu_ex_ in the “up” conformation. A subsequent H^+^-induced conformational change generates a state D (captured in the QQQ mutant structure), which has an open extracellular vestibule and new positionings for Glu_ex_ and Glu_in_. Conformational dynamics of Glu_in_ allows water pathways to connect Glu_ex_ directly to the intracellular bulk water, as in state E. From State E, deprotonation of Glu_ex_ promotes its return to the anion pathway, in competition for Cl^−^ (state F). Binding of Cl^−^ from the intracellular side, coordinated with inner-gate opening (Basilio et al., 2014) (reflected by the dotted lines) generates the original state A. (**B**) The updated transport model is consistent with Cl^−^/H^+^ exchange seen in transporters that do not have a titratable residue at the Glu_in_ position, such as cmCLC. The left panel shows a structural overlay of QQQ and cmCLC, highlighting the positions of the Thr and Lys residues occurring at the Glu_in_ and E113 positions respectively. The right panel is a cartoon depiction of the water pathway. (**C**) The CLC-1 channel structure shows a Glu_ex_ “out” position analogous to that seen in the QQQ mutant. Overlays of CLC-ec1 WT (grey), QQQ (purple), and CLC- 1 (green) illustrate changes in positioning of conserved residues near Glu_ex_.

State D involves a widening of the extracellular vestibule, which will facilitate Cl^−^ binding from and release to the extracellular side. In the QQQ structure (our approximation of State D), the reorientation of Helix N results in subtle changes in Cl^−^-coordination at the S_ext_ site (**Figure 3 – figure supplement 2**), which suggests that binding at this site may be weakened, though we currently lack direct evidence for this conjecture. Regardless of the affinity at S_ext_, the opening of the extracellular permeation pathway in State D will promote Cl^−^ exchange in both directions, which is essential to achieving reversible transport.

In addition to involving a widening of the extracellular vestibule, state D has the protonated Glu_ex_ in an “out” conformation and within ∼5 Å of Glu_in_ (**Figure 1C,D**). At first glance, this positioning suggested to us that Glu_in_ might be participating in an almost direct hand-off of H^+^ to and from Glu_ex_. However, MD simulations revealed that Glu_in_ is highly dynamic and most often is rotated away from its starting position, allowing the robust formation of water pathways from the intracellular bulk water directly to Glu_ex_ (**Figure 5**) (State E). Once such transfer occurs, the deprotonated Glu_ex_ will be disfavored in the hydrophobic core, and it will compete with Cl^−^ for the S_cen_ anion-binding site, generating State F. Although this conformational state has not been observed crystallographically for CLC-ec1, the computational studies of Piccolo et al. (Picollo et al., 2012) found that Glu_ex_ favors the S_cen_ position when there are no Cl^−^ ions bound in the pathway (as in State F), and those of Mayes et al. found that the “down” position is in general the preferred orientation for Glu_ex_ (Mayes et al., 2018). In addition, a recent structure of an Asp_ex_ CLC-ec1 mutant supports that the carboxylate likes to reach down towards S_cen_, in a “midlow” position, excluding the presence of Cl^−^ at both S_cen_ and S_ext_ (Park et al., 2019), as depicted in State F. From this state, binding of Cl^−^ from the intracellular side (coordinated with inner-gate opening (Basilio et al., 2014)) knocks Glu_ex_ back up to S_ext_, generating the original state A. This transport cycle is fully reversible, allowing efficient transport in both directions, as is observed experimentally (Matulef and Maduke, 2005).

## DISCUSSION

In this study, we aspired to determine the high-resolution structure of the CLC “outward-facing open” conformational state. Our conclusion that the QQQ mutant structure represents such an intermediate in the CLC transport mechanism is supported by several pieces of evidence. First, the WT protein under low-pH conditions (glutamate residues protonated), adopts a conformation different from the high-pH condition, as detected by DEER spectroscopy, with the conformational change in the direction predicted by the QQQ structure (**Figure 4A-D**). Second, the movement of helix P in the QQQ structure is as predicted by the fact that cross-linking helix P inhibits transport (Khantwal et al., 2016) (**Figure 4E**). Third, and perhaps most compellingly, the details observed in this conformational state reconcile a multitude of findings in the literature.

Our proposed updated transport model (**Figure 7A**), in addition to retaining key features based on previous models (Miller and Nguitragool, 2009; Feng et al., 2012; Basilio et al., 2014; Khantwal et al., 2016), unifies our picture of both CLC transporter and channel mechanisms. First, it is compatible with transporters that have non-titratable residues at Glu_in_ and E113. Our simulations and experiments (**Figures 5 and 6**) lead to the conclusion that these residues play a key role in regulating water pathways rather than in direct hand-off of H^+^. From this perspective, the evolution of non-titratable residues in either (Feng et al., 2010) or both (Phillips et al., 2012; Stockbridge et al., 2012) of these positions is perfectly sensible. In addition, previous mutagenesis experiments on CLC-ec1, which demonstrated a surprising tolerance for mutations at Glu_in_ (Lim and Miller, 2009), now make more sense. Strikingly, the structural positioning of T269 in cmCLC, located at the Glu_in_ sequence position, matches the structural positioning of Gln_in_ in the QQQ mutant, such that side chain dynamics could facilitate comparable water pathways (**Figure 7B**).

The second unifying feature of our model is that it attests to Glu_ex_ movements being conserved amongst every known type of CLC: 2:1 Cl^−^/H^+^ exchangers, 1:1 F^−^/H^+^ exchangers, and uncoupled Cl^−^ channels. Previously, an “out” position for Glu_ex_ had been proposed to be essential to the mechanism of F^−^/H^+^ exchangers (Last et al., 2018), which allow bacteria to resist fluoride toxicity (Stockbridge et al., 2012). However, such a conformation had not been directly observed, and it was postulated that it may be only relevant to the F^−^/H^+^ branch of the CLC family. Structurally, Glu_ex_ in the “out” position has previously only been observed in a CLC channel. This structure appears to represent an open-channel conformation, and thus was deduced that the “out” position is relevant only to CLC channels, particularly because of residue clashes that were predicted based on transporter structures known at the time (Park and MacKinnon, 2018). An overlay of the CLC-ec1 QQQ structure with the hCLC-1 channel structure reveals that the Glu_ex_ “out” conformations are similar (**Figure 7C**). Thus, this conformation is a unifying feature of CLC channels and transporters. Moreover, this conclusion connotes that all CLC proteins act via a “windmill” mechanism (Last et al., 2018), in which the protonated Glu_ex_ favors the core of the protein while the deprotonated Glu_ex_ favors the anion-permeation pathway. Such a mechanism is preferable to previous “piston”-type mechanisms, with Glu_ex_ moving up and down within the anion-permeation pathway, which required a protonated (neutral) Glu_ex_ to compete with negatively charged Cl^−^ ions.

Elements of the transport cycle require future experiments to elucidate details. Prominently, the nature of the inward-facing conformational state remains uncertain. In our model, we indicated inward-opening with dotted lines (**Figure 7**, States F, A, B) to reflect this uncertainty. One proposal is that the inner-gate area remains static and transport works via a kinetic barrier to Cl^−^ movement to and from the intracellular side (Feng et al., 2010). Consistent with this proposal, multiscale kinetic modeling revealed that 2:1 Cl^−^/H^+^ exchange can arise from kinetic coupling alone, without the need for large protein conformational change (Mayes et al., 2018). An alternative proposal is that CLCs visit a conformationally distinct inward-open state, based on the finding that transport activity is inhibited by cross-links that restrict motion of Helix O, located adjacent to the inner gate (Basilio et al., 2014; Accardi, 2015). This putative inward-open state appears distinct from the conformational change observed in the QQQ mutant, as the inter-residue distances for the cross-link pairs (399/432 and 399/259) are unchanged in QQQ relative to WT. The details of the kinetic-barrier and conformational-change models, and the need for additional experiments on this aspect of transport, have been clearly and comprehensively discussed (Accardi, 2015; Jentsch and Pusch, 2018).

Despite having an open extracellular vestibule, the QQQ mutant is a slow transporter – almost an order of magnitude slower than the wild-type protein (at pH 4.5). This relative slowness may be due to the QQQ’s increased Cl^−^ binding affinity (140 µM, **Figure 3 – Figure supplement 2**, vs 600 µM for WT (Picollo et al., 2009; Khantwal et al., 2016)) and hence slower dissociation rate (Picollo et al., 2009). Increased Cl^−^-binding affinity (relative to WT) is observed in all Glu_ex_ mutants (Picollo et al., 2009; Picollo et al., 2012; Park et al., 2019), likely because no carboxylate side chain is available to compete Cl^−^ out of the binding sites.

Simulations of the QQQ conformational state with the glutamine residues reverted to the native, protonatable glutamate side chains will be needed for full understanding of how protonation and deprotonation of these residues affect the conformational dynamics of side chains and water pathways. In our current model, we propose that Glu_in_ needs to be in the protonated (neutral) state to adopt the position that allows water pathways. This proposal appears harmonious with the hydrophobic nature of the protein core explored by the Glu_in_ side chain and the fact that other CLC homologs use neutral residues at this position (Feng et al., 2010; Phillips et al., 2012; Stockbridge et al., 2012). In addition, the proposal is consistent with MD simulations that show the Glu_ex_/Glu_in_ doubly protonated state is highly populated (Mayes et al., 2018) and can favor formation of water pathways under certain conditions (Ko and Jo, 2010). Nevertheless, simulations with glutamate side chains in the QQQ conformational state, together with explicit evaluation of H^+^ transport (Wang et al., 2018; Duster et al., 2019), are needed to elaborate details of the H^+^-transfer steps. In addition, multiscale modeling can expand the picture to include multiple pathways that are likely to occur (Mayes et al., 2018). Recognizing the importance of elaborating these details, the results reported here represent an essential and pivotal step toward a complete, molecularly detailed description of mechanism in the *sui generis* CLC transporters and channels.

## MATERIALS AND METHODS

### Protein preparation and purification

Mutations were inserted in the wild type CLC-ec1 protein using Agilent QuikChange Lightning kit and were confirmed by sequencing. Protein purification was carried out as described (Walden et al., 2007), with a few changes to the protocol for ITC and crystallization experiments. For ITC, QQQ or E148Q were purified in buffer A (150mM Na-isethionate, 10mM HEPES, 5mM anagrade decyl maltoside (DM) at pH 7.5). For crystallization experiments, QQQ was extracted with DM. The detergent was gradually exchanged for lauryl maltose neopentyl glycol (LMNG) during the cobalt-affinity chromatography step. The final size-exclusion chromatography step was performed in a buffer containing LMNG. All detergents were purchased from Anatrace (Maumee, OH).

### Crystallography

Purified QQQ protein was concentrated to at least 30 mg/mL. Concentrated protein was mixed with 1.5 parts (w/w) of monolein containing 10% (w/w) cholesterol using the syringe reconstitution method (Caffrey and Cherezov, 2009), to generate a lipidic cubic phase mixture. 25-nL droplets of the mixture was dispensed on glass plates and overlaid with 600 nL of precipitant using a Gryphon crystallization robot (Art Robbins Instruments, Sunnyvale, CA). Crystallization trials were performed in 96-well glass sandwich plates incubated at 16 °C. The best crystals were obtained using a precipitant solution consisting of 100mM Tris (pH 8.5), 100 mM sodium malonate, 30% PEG 400 and 2.5% MPD Crystals were harvested after 3-4 weeks of incubation and flash-frozen in liquid nitrogen without further additives.

### Structure determination and refinement

X-ray diffraction data were collected at APS at GM/CA beamline 23ID-D and were processed using XDS (Kabsch, 2010) and AIMLESS (Evans, 2006) from the CCP4 suite (Winn et al., 2011). Owing to radiation damage, a complete dataset was collected by merging data from 3 different crystals. Phases were obtained using PHASER (McCoy et al., 2007) with PDB ID 1ots as a search model. Iterative refinement was performed manually in Coot (Emsley and Cowtan, 2004) and REFMAC (Murshudov et al., 1997). The final model contained all residues except those of helix A due to lack of density for this region of the protein. Helix A is observed in different conformations in the monomeric versus dimeric CLC-ec1 structures, and has no impact on function (Robertson et al., 2010).

### Reconstitution and chloride flux assay

Flux assay results presented in this paper required a variety of experimental conditions for reconstitutions and flux assays, summarized in **Table 2**. For flux assays comparing activity at pH 7.5 and 4.5, purified CLC-ec1 were first reconstituted at pH 6. The samples were then aliquoted and pH-adjusted using a 9:1 ratio of sample and the adjustment buffer. This step was taken to eliminate variability from separate reconstitutions. For experiments testing H^+^ pumping in mutants, a pH gradient was used to ensure any measured H^+^ transport was from H^+^ pumping and not H^+^ leak.

**Table 2.** Buffers used for Reconstitution and Flux Assays.

| Buffer R | Buffer F | Ion flux monitored | Protein-lipid ratio (µg/mg) | Protein per assay (µg) |
| --- | --- | --- | --- | --- |
| <u>Comparing turnover rates at pH 7.5 and 4.5 (Figure 2)</u> |  |  |  |  |
| 333 mM KCl, 55 mM Na-citrate, 55 mM Na<sub>2</sub>HPO<sub>4</sub>, pH 6.0 | 333 mM K-isethionate, 55 µM KCl, 55 mM Na-citrate, 55 mM Na<sub>2</sub>HPO<sub>4</sub>, pH 6.0 | Cl<sup>-</sup> | 0.4 - 0.8 | 0.4 - 0.8 |
| pH adjustments by adjustment buffers (10x) (after reconstitution): | pH adjustments by adjustment buffers (10x): |  |  |  |
| 0.16 M citric acid, 0.42 M Na<sub>3</sub>PO<sub>4</sub> (for pH 7.5)<br>0.43 M citric acid, 0.29 M Na<sub>3</sub>PO<sub>4</sub> (for pH 4.5) | 0.16 M citric acid, 0.42 M Na<sub>3</sub>PO<sub>4</sub> (for pH 7.5)<br>0.43 M citric acid, 0.29 M Na<sub>3</sub>PO<sub>4</sub> (for pH 4.5) |  |  |  |
| <u>Testing H<sup>+</sup> pumping of mutants (Figure 6)</u> |  |  |  |  |
| 300 mM KCl, 40 mM Na-citrate, pH 4.0 | 300 mM K-isethionate, 50 µM KCl, 2 mM Na-citrate, pH 4.3 | H<sup>+</sup> and Cl<sup>-</sup> | 0.4 - 5.0 | 0.4 - 10 |
| <u>Testing DEER samples (Figure 4 – figure supplement 1)</u> |  |  |  |  |
| 300 mM KCl, 40 mM Na-citrate, pH 4.5 | 300 mM K-isethionate, 50 µM KCl, 2 mM Na-citrate, pH 4.5 | Cl<sup>-</sup> | 0.4 | 0.4 - 0.5 |

To measure the rate of H^+^ and Cl^−^ transport in flux assays, purified CLC-ec1 proteins were reconstituted into phospholipid vesicles (Walden et al, 2007). *E coli* polar lipids (Avanti Polar Lipids, Alabaster, AL) in chloroform were dried under argon in a round-bottomed flask. To ensure complete removal of chloroform, the lipids were subsequently dissolved in pentane and dried under vacuum on a rotator, followed by further drying (5 minutes) under argon. The lipids were then solubilized at 20 mg/mL in buffer R (**Table 2**) with 35 mM CHAPS on the rotator for 1.5-2 hours. Purified proteins (0.4 - 5 µg per mg lipids) were added to the prepared lipid-detergent mix. The detergent in the samples was then gradually removed by dialysis over 2 nights. Each reconstitution sample was divided into two for measurement in duplicate. Duplicates were averaged to obtain a turnover rate value. (In reporting results of experiments, each “n” is an average of duplicates from a reconstitution sample.)

Reconstituted vesicles were subjected to 4 freeze-thaw cycles and were then extruded with an Avanti Mini Extruder using a 0.4 µm-filter (GE Healthcare, Chicago, IL) 15 times. For each assay, 60 - 120 µL of extruded sample were buffer-exchanged through 1.5- to 3.0-mL Sephadex G-50 Fine resin (GE Healthcare, Chicago, IL) columns equilibrated with buffer F (**Table 2**). Exchange was accomplished by spinning the columns at ∼1100 g for 90 seconds using a clinical centrifuge. The collected sample (80 - 200 µL) was then added to buffer F (500 - 600 µL) for flux-assay measurement. Extravesicular [Cl^−^] and [H^+^] were monitored using a Ag·AgCl electrode and a pH electrode, respectively. The electrodes were calibrated by known additions of KCl (in 20-136 nmol steps) and NaOH (in 10-50 nmol steps). Sustained ion transport by CLC-ec1 was initiated by addition of 1.7-3.4 µg/mL of valinomycin (from 0.5 mg/mL stock solution in ethanol).

### Isothermal titration calorimetry

Titration isotherms were obtained using a VP-ITC microcalorimeter (MicroCal LLC, Northampton, MA) at 25 °C. For the experiment, QQQ or E148Q protein were purified in buffer A. Titrant used in the experiment was 30 mM KCl in buffer A. The starting concentration of protein was 15-20 µM, in a volume of 1.5 mL. KCl (30 mM) was syringe-titrated into the sample cell in thirty 10-µL injections. The reference data were obtained by titrating buffer A into the protein-containing solution. Data were analyzed using Origin 7.0 software, with fitting using the “one set of sites” model (keeping n=1). The other thermodynamic parameters were obtained accordingly. Isethionate was chosen as the anion of choice for purification of proteins for the ITC experiments since the QQQ mutant shows aggregation upon purification in tartrate-containing solutions, which were previously used (Picollo et al., 2009; Khantwal et al., 2016). The mutant is comparatively stable in isethionate and continues to remain stable throughout the ITC experiment. Figure 3 – figure supplement 2 shows the gel filtration chromatograms of the mutants before and after the ITC experiments.

### DEER spectroscopy

Protein purification and sample preparation for proteins used in DEER spectroscopy was performed as described (Khantwal et al., 2016). DEER experiments were performed at 83 K on a Bruker 580 pulsed EPR spectrometer at Q-band frequency (33.5 GHz) using either a standard four-pulse protocol (Jeschke and Polyhach, 2007) or a five-pulse protocol (Borbat et al., 2013). Analysis of the DEER data to determine P(r) distance distributions was carried out in homemade software running in MATLAB (Brandon et al., 2012; Stein et al., 2015). In the original five-pulse protocol paper the pure five-pulse signal was obtained by subtracting the artefact four-pulse data (Borbat et al., 2013). This method requires the ability to discern clearly the extent of the artefact. For the data in this study, we chose to simultaneously fit the four- and five-pulse data with a single Gaussian component in order to improve accuracy of subtracting the four-pulse artefact (**Figure 4 – figure supplement 2**). Confidence bands for the distance distributions were determined using the delta method (Hustedt et al., 2018). The confidence bands define the 95% confidence interval that the best fit distance distribution will have. In the case of a Gaussian distribution, the shape of the confidence bands can be non-Gaussian.

### Simulation system setup

The structure of the CLC-ec1 QQQ mutant crystallized in this work at 2.6-Å resolution was used as the starting structure for the MD simulation. The 2 Cl^−^ ions bound at S_cen_ and S_int_ sites in each of the two subunits were preserved for the simulation. In our initial refinement of the QQQ structure, we had modeled water rather than Cl^−^ at the S_ext_ site, and therefore the simulation was performed without Cl^−^ at this site. The pKa of each ionizable residue was estimated using PROPKA (Olsson et al., 2011; Rostkowski et al., 2011), and the protonation states were assigned based on the pKa analysis at pH 4.5. Missing hydrogen atoms were added using PSFGEN in VMD (Humphrey et al., 1996). In addition to the crystallographically resolved water molecules, internal water molecules were placed in energetically favorable positions within the protein using DOWSER (Zhang and Hermans, 1996; Morozenko et al., 2014). One of the energetically favorable water molecules was added right between the side chains of Q148 and Q203, nicely bridging the two residues. The QQQ protein was embedded in a POPE lipid bilayer using the CHARMM-GUI membrane builder (Wu et al., 2014). The membrane/protein system was fully solvated with TIP3P water (Jorgensen et al., 1983) and buffered in 150 mM NaCl to keep the system neutral. The resulting systems consisting of ∼155,000 atoms were contained in a 164 × 127 × 98-Å^3^ simulation box.

### Simulation protocols

MD simulation was carried out with NAMD2.12 (Phillips et al., 2005) using CHARMM36 force field (Klauda et al., 2010; Huang and MacKerell, 2013) and a time step of 2 fs. Periodic boundary conditions were used throughout the simulations. To evaluate long-range electrostatic interactions without truncation, the particle mesh Ewald method (Darden et al., 1998) was used. A smoothing function was employed for short-range nonbonded van der Waals forces starting at a distance of 10 Å with a cutoff of 12 Å. Bonded interactions and short-range nonbonded interactions were calculated every 2 fs. Pairs of atoms whose interactions were evaluated were searched and updated every 20 fs. A cutoff (13.5 Å) slightly longer than the nonbonded cutoff was applied to search for interacting atom pairs. Simulation systems were subjected to Langevin dynamics and the Nosé–Hoover Langevin piston method (Nose, 1984; Hoover, 1985) to maintain constant pressure (P = 1 atm) and temperature (T = 310 K) (NPT ensemble).

The simulation system was energy-minimized for 10,000 steps, followed by two stages of 1-ns relaxation. Both the protein and the Cl^−^ ions in the binding sites were positionally restrained (k = 1 kcal·mol^−1^·Å^−2^) in the first 1-ns simulation to allow the membrane to relax. In the second 1-ns simulation, only the protein backbone and the bound Cl^−^ ions were positionally restrained (k = 1 kcal·mol^−1^·Å^−2^) to allow the protein side chains to relax. Then a 600-ns equilibrium simulation was performed for the system without any restraint applied.

### Analysis of water pathways

The water pathways between Q148 (Gln_ex_) and the intracellular bulk water was searched using a breadth-first algorithm. In subunit 2 of the homodimer, Gln_ex_ drifted up and away from the “out” position at the beginning of the simulation (within 5 ns), and it did not return to the “out” position during the simulation. In subunit 1, Gln_ex_ remained near the “out” position for the first 400 ns; we focused our analysis of water pathways on this subunit. The simple distance criterion of 2.5-Å for the hydrogen bonds, which was found to be very useful and cheapest in computational terms in a previous study (Matsumoto, 2007) was used to determine whether water molecules are connected through continuous hydrogen-bonded network. The water pathway with the smallest number of O-H bonds in each frame was considered as the shortest path. The first water molecule in each water pathway is searched using a distance cutoff of 3.5-Å for any water oxygen atoms near the OE1/NE2 atoms of Q148. The water pathway is considered to reach the intracellular bulk once the oxygen atom of the newly found water molecules is at z < −15 Å (membrane center is at z = 0).

## ACKNOWLEDGMENTS

We thank Chris Miller and Martin Prieto for comments on the manuscript. We are grateful to Brian Kobilka for use of the crystallization equipment and to K. Chris Garcia for use of the MicroCal ITC instrument. This research was funded by NIH GM113195 (M.M., E.T., and H.S.M.). T.S.C. was support by an American Heart Association Fellowship 17POST33670553. This research used resources of the Advanced Photon Source, a US Department of Energy (DOE) Office of Science User Facility operated for the DOE Office of Science by Argonne National Laboratory under contract no. DE-AC02-06CH11357. The SSRL Structural Molecular Biology Program is supported by the DOE Office of Biological and Environmental Research, and by the National Institutes of Health, National Institute of General Medical Sciences (P41GM103393). We also acknowledge computing resources provided by Blue Waters at National Center for Supercomputing Applications (T.J.), and Extreme Science and Engineering Discovery Environment (grant MCA06N060 to E.T.).

---

**Figure 3 – Figure supplement 1.**
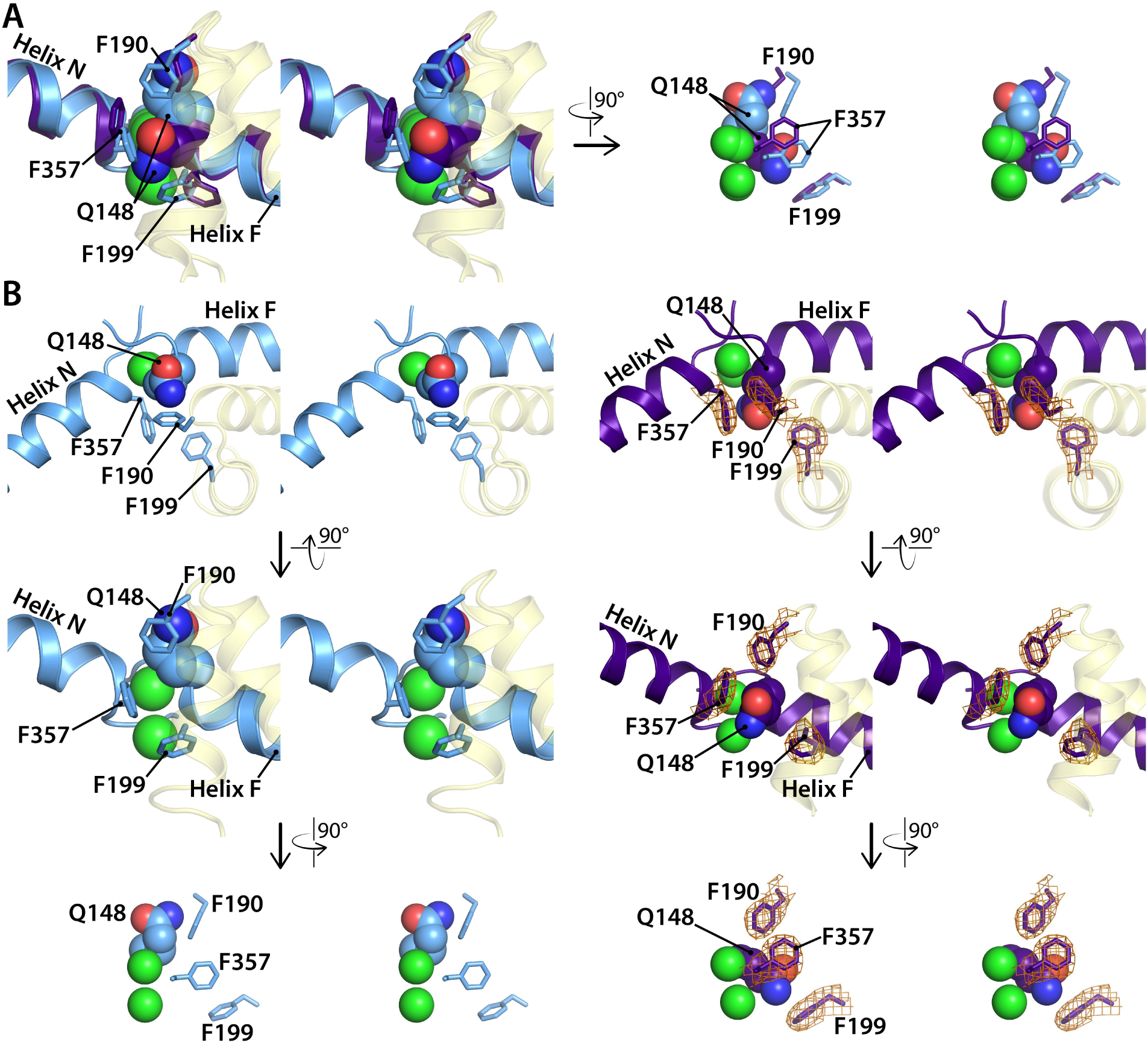
Comparison of the CLC-ec1 QQQ structure to the E148Q (Glu_ex_ to Gln) structure in the vicinity of the S_ext_ binding site. All diagrams are in stereoview. (**A**) Overlay of E148Q (blue) and QQQ (purple). Conserved residues F190, F199, and F357 are shown as sticks. (**B**) Side-by-side comparison of E148Q (left) and QQQ (right), with electron density shown (2Fo-Fc map at 1σ contour level) in QQQ for the repositioned residues F190, F199, and F357.

**Figure 3 – figure supplement 2.**
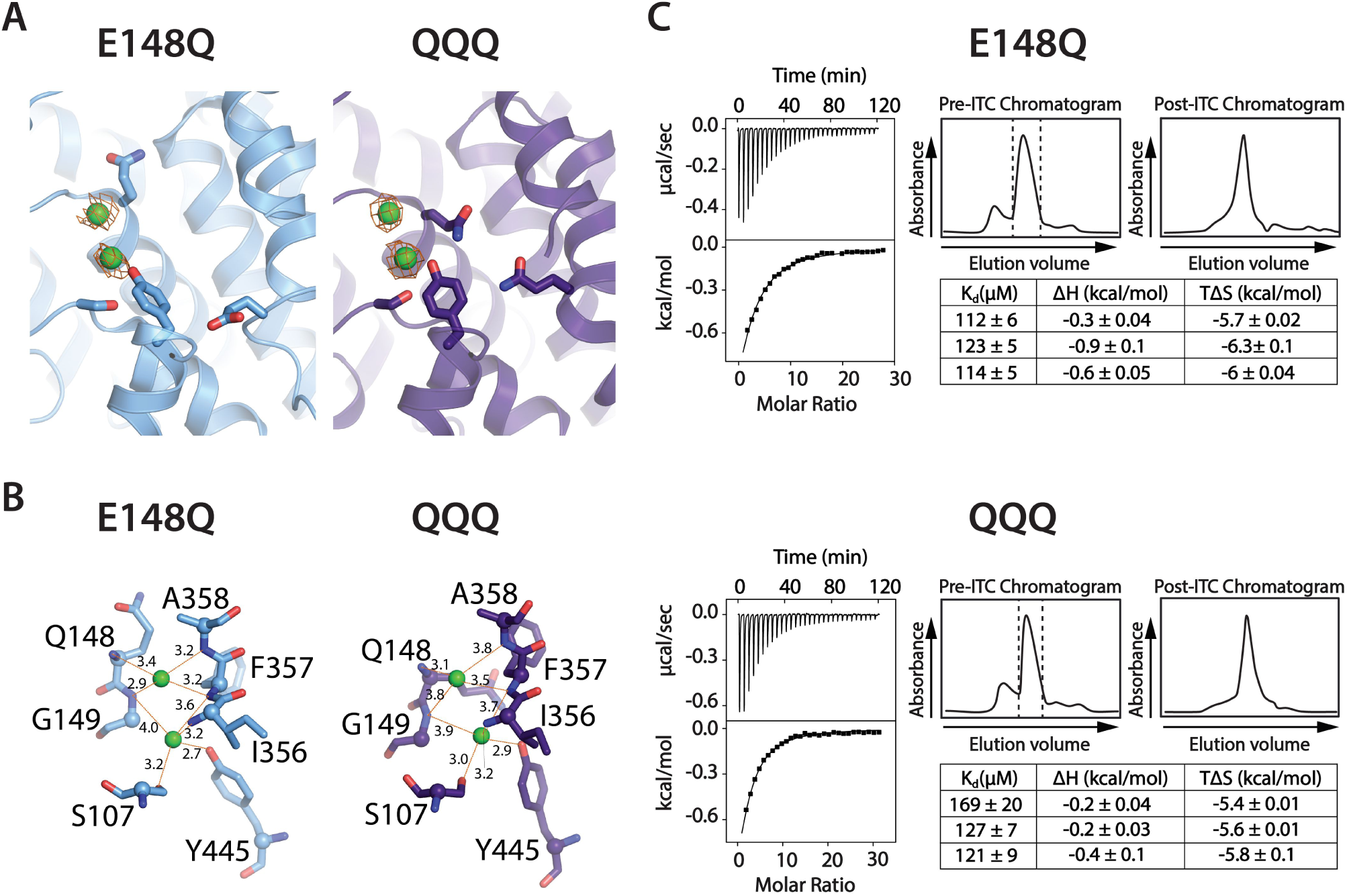
Cl^−^ binding to QQQ CLC-ec1. (**A**) 2Fo-Fc maps (1σ contour level) for QQQ and E148Q show electron density consistent with Cl^−^ binding to both the S_cen_ and S_ext_ sites. (**B**) Close-up views of the S_ext_ site in E148Q and QQQ, illustrating the increased protein/Cl^−^ coordination distances in the QQQ mutant. The coordination distances at the S_cen_ site are identical between E148Q and QQQ. **(C)** Cl^−^ binding measured by ITC for E148Q (top) and QQQ (bottom). At left are shown representative data from ITC experiments. At right are shown representative size-exclusion chromatograms of samples before and after the ITC experiment (pre- and post-ITC chromatograms). The dotted line in the pre-ITC chromatogram represents the fraction that was collected for use in the ITC experiment. The sample was run again following the ITC experiment to confirm the sample was stable throughout the experiment. The summary tables show results for 3 experiments performed on 3 separate protein preparations. Averages (± SEM) are as follows: for E148Q, K_d_ = 116 ± 6 µM, ΔH = −0.6 ± 0.3 kcal/mol, TΔS = −6 ± 0.3 kcal/mol; for QQQ, K_d_, = 138 ± 26 µM, ΔH = −0.2 ± 0.1 kcal/mol, TΔS = −5.6 ± 0.2 kcal/mol. These measurements do not distinguish between binding to S_cen_ versus binding to S_ext_. Previously, crystallographic experiments were used to distinguish binding at the different sites (Lobet and Dutzler, 2006). We attempted to perform such experiments on QQQ; however, these experiments make use of the anomalous signal obtained from Br^−^ binding, and we have not been successful in our attempts to obtain quality diffracting QQQ crystals in the presence of Br^−^.

**Figure 3 – figure supplement 3.**
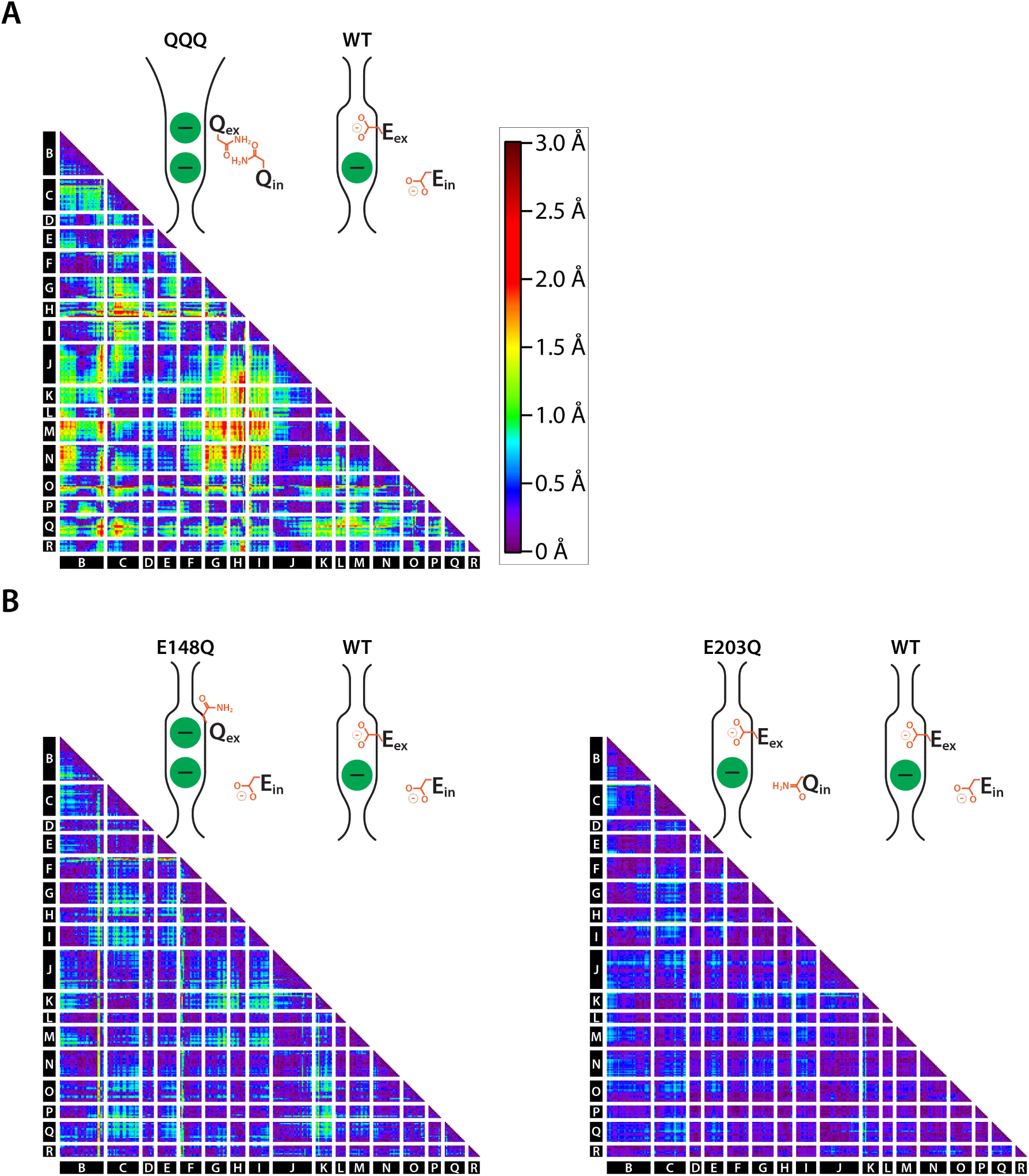
Cα difference distances matrices. (**A**) Difference distance matrix comparing WT CLC-ec1 (1ots) to QQQ reveals intramolecular rearrangements throughout the protein, most notably at Helices C, G, H, L, M, N, and Q. (**B**) Comparison of WT to E148Q (1otu) or E203Q (2fec) reveals only minor (≤ 0.8 Å) changes.

**Figure 3 – figure supplement 4.**
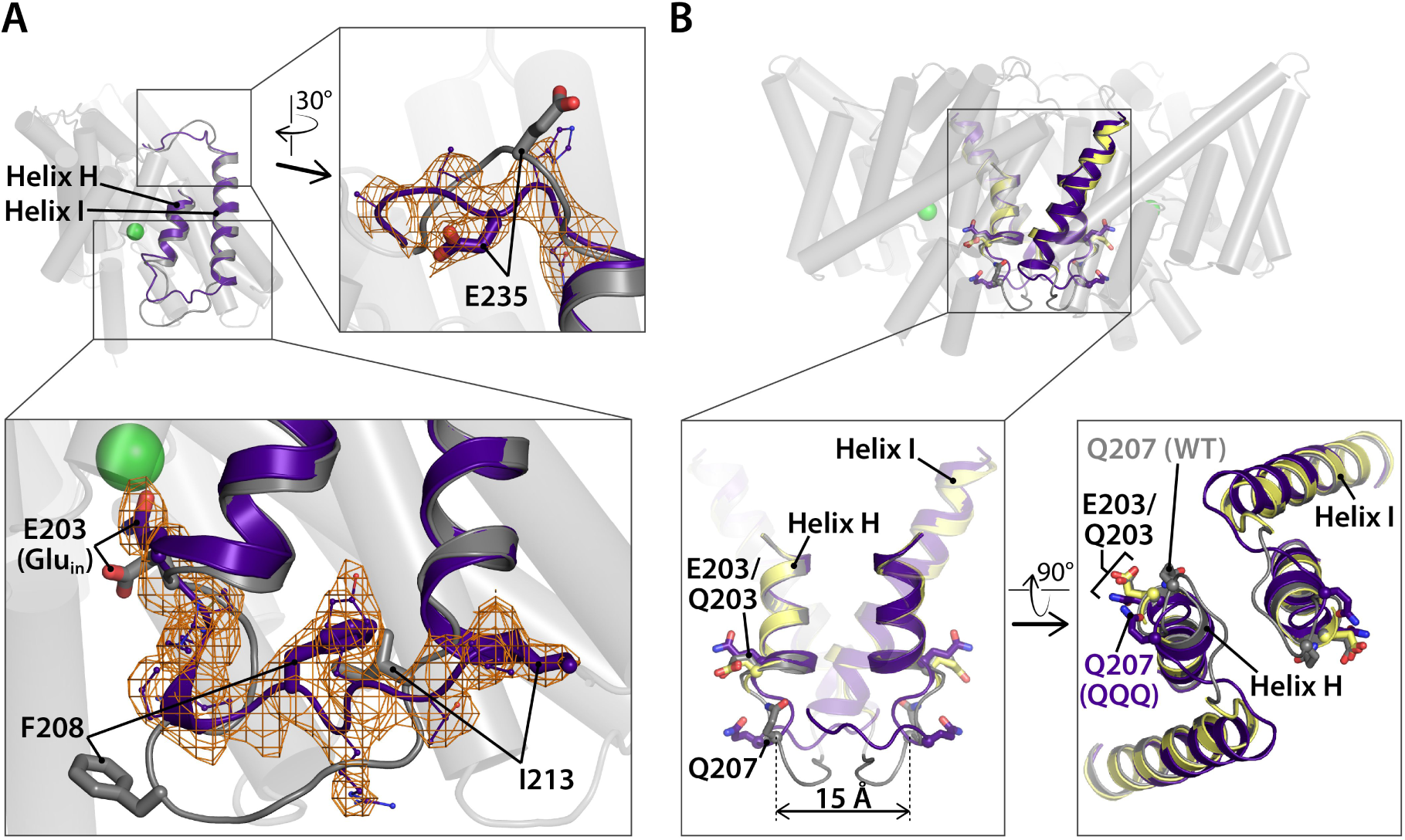
Linker rearrangements in the QQQ structure. (**A**) Structural overlay of WT CLC-ec1 (1ots) and QQQ. The WT structure (one subunit) is shown in grey. Overlaid in purple is the structure of QQQ from residues 203 to 237, encompassing Helices H and I in addition to the H-I and I-J linkers, which are displaced in QQQ compared to WT. Electron density is shown for the HI linker (residues 205 to 213). (**B**) Views of the H-I linker in the context of previous cross-linking results. In WT (grey) and QQQ (purple), residues Q207 on the two subunits are separated by 15 Å. In the Q207C structure (yellow), in which the Q207C residues are cross-linked, the H-I linker is not visible, and thus the exact positioning of the disulfide bond is uncertain; however, the observation that the H-I helices (including E203, Glu_in_) are unaffected by the cross-link illustrates that H-I linker movement does not affect the main body of the protein.

**Figure 4 – figure supplement 1.**
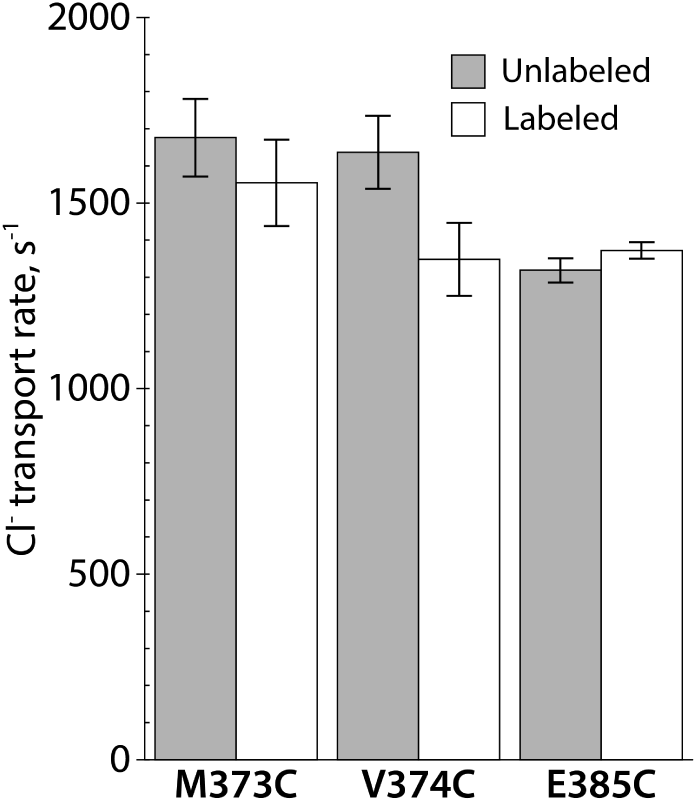
Activity of spin-labeled DEER samples. Protein was reconstituted into liposomes before and after spin-labeling at the cysteine residue. Unitary Cl^−^ transport rates are shown, ± SEM, n= 2 for each sample.

**Figure 4 – figure supplement 2.**
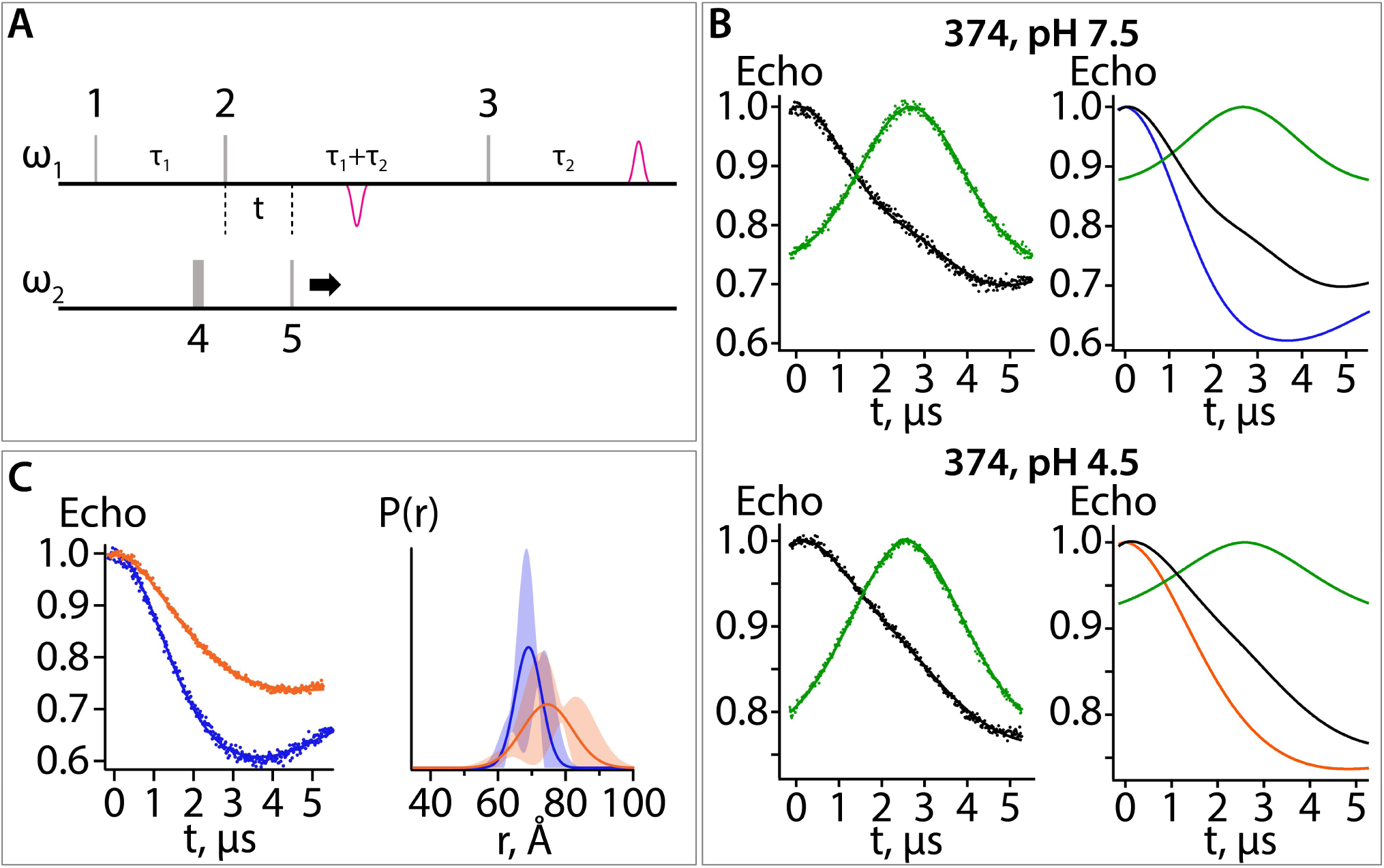
5-pulse DEER experiment: breakdown of the fitting procedure. (**A**) Diagram of the 5-pulse DEER experiment. Pulses 1-3 are at frequency 1 (ω_1_) and pulses 4 and 5 are at frequency 2 (ω_2_). Pulses 1-3 are square pulses of length 16, 32, and 32 ns, respectively. Pulse 5 is a 200 ns chirp pulse starting at 77 MHz and ending at 47 MHz from frequency 1. Pulse 4 is a 40 ns square pulse positioned 62 MHz from frequency 1. The time intervals, τ_1_ and τ_2_, are 2.8 and 2.9 μs, respectively. (**B**) Data for sample with spin label at position 374 at pH 7.5 (upper panels) and 4.5 (lower panels). The data shown in the left panels were collected with all 5 pulses (black) or with the amplitude of pulse 5 set to zero (green). The former experiment provides a combination of a pure five-pulse decay and a four-pulse artefact; the latter provides a measure of the four-pulse artefact. These data were analyzed to obtain the pure 5-pulse traces, shown as the blue and orange traces in the right panels. The 5-pulse experimental results (black) are a scaled sum of the pure 5-pulse traces (blue for pH 7.5; orange for pH 4.5) and the scaled 4-pulse artefact (green). The scaling factors are obtained as variables in the fitting routine. (**C**) On the left are the pure five-pulse data obtained from removing the four-pulse artefact, as described in (B). On the right are the single Gaussian distributions and confidence bands for the two conditions. These data are the same as those shown in Fig. 4C.

**Figure 5 – figure supplement 1.**
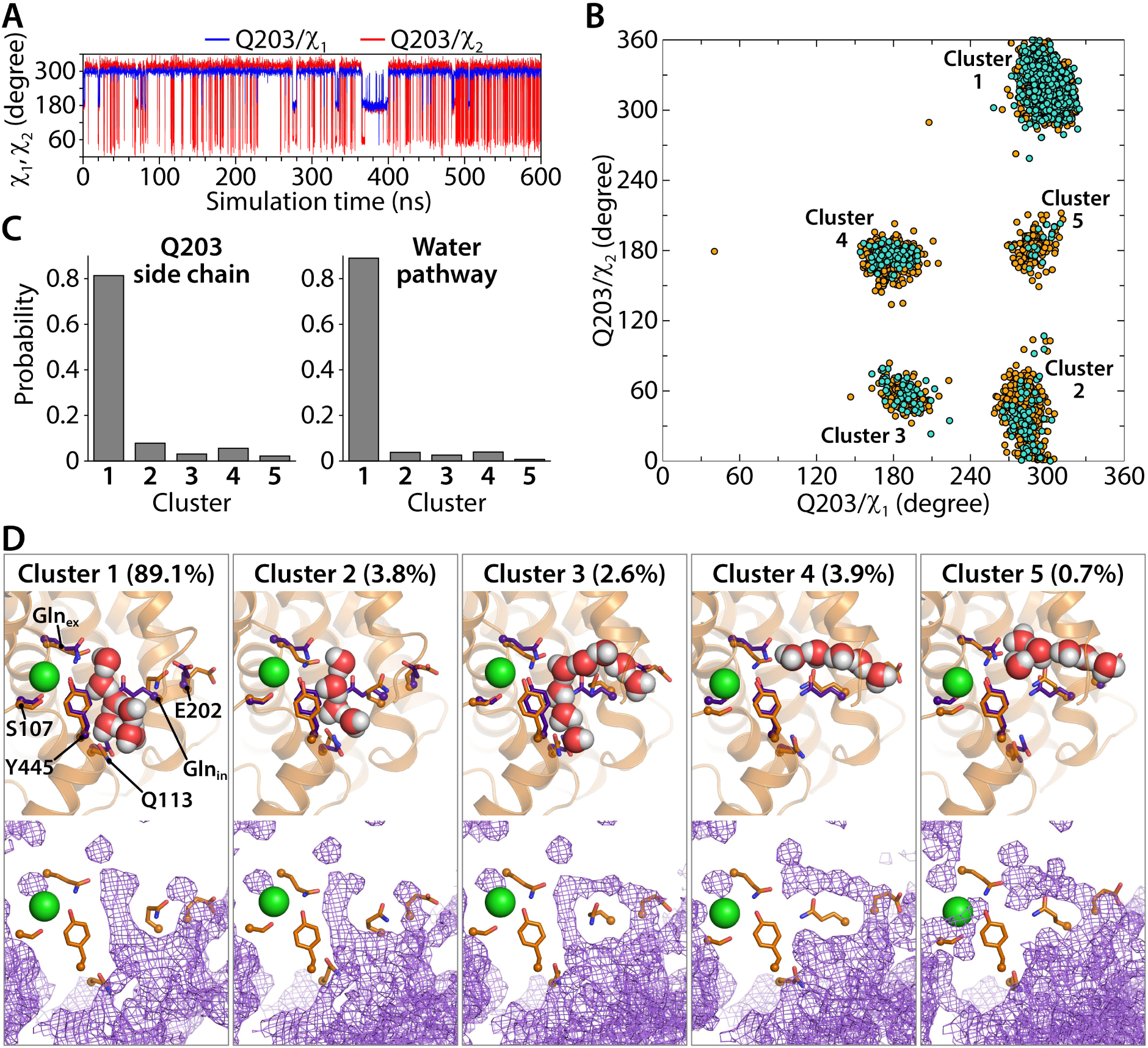
Water pathways and conformational dynamics of Gln_in_ (Q203) side chain orientation. (**A**) Traced dihedral angles of Q203 during the simulation of the CLC-ec1 QQQ mutant show that the Q203 side chain is flexible and undergoes dynamical changes over the course of the simulation. (**B**) 2D scatter map of Q203 dihedral angles reveals 5 main clusters of side chain conformations sampled during the equilibrium simulation. Each dot represents one pair of the dihedral angles of Q203 from snapshots taken at 100-ps intervals in the simulation. The frames with continuous water pathways are shown in cyan. (**C**) The distribution of the side chain conformations of Q203 over the trajectory (left panel) shows that Cluster 1 is most populated. This conformation also captures most of the water pathways (right panel) during the simulation. The water-pathway probability is normalized to the total number of water pathways. (**D**) Representative simulation snapshots with Gln_in_ in different conformations, showing continuous water pathways between Gln_ex_ and the intracellular bulk. Clusters 4 and 5, representing less than 5% of the pathways, reach intracellular bulk water via an alternate pathway. Lower panels show the overall water occupancy maps for each cluster in the simulation. Density is contoured at isovalue 0.35.

**Figure 5 – figure supplement 2.**
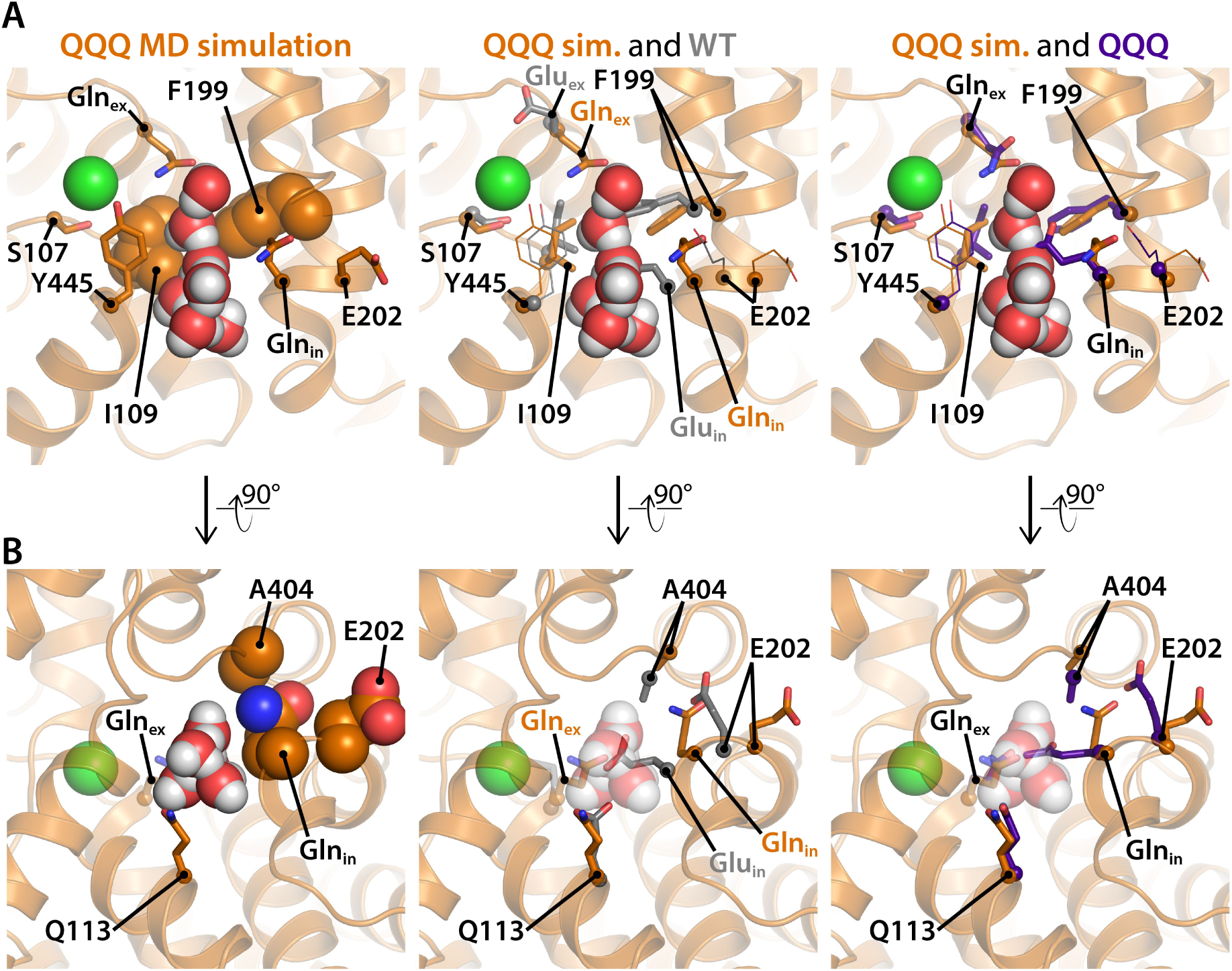
Residues of interest near the inner water pathway. (**A**) The left panel shows that the inner water pathway is lined by F199 and I109; at the latter position, mutations were found to specifically inhibit the H^+^ branch of the CLC transport cycle (Han et al., 2014). S107 and Y445, located at the inner gate of the Cl^−^-permeation pathway, are shown for orientation. The middle and right panels show overlays to compare side chain positioning in the simulation snapshot to side chain position in the WT and QQQ crystal structures, respectively. Gln_in_ and F199 side chains project into the water pathway and must rearrange to accommodate water entry into the cavity. (**B**) A rotated view of the inner water pathway shows tight packing of residues around Glu_in_/Gln_in_ and the “interfacial pathway” residues E202, and A404 (left panel). The same view with overlay of side chains from the WT crystal structure (middle panel) and the QQQ crystal structure (right panel) show that water entry requires rotation of Glu_in_/Gln_in_ away from crystallographic conformations. The tight packing explains how substitutions of large residues for E202 or A404 (Lim et al., 2012; Han et al., 2014) would obstruct Glu_in_/Gln_in_ from moving to accommodate the inner water pathway.

**Figure 5 – figure supplement 3.**
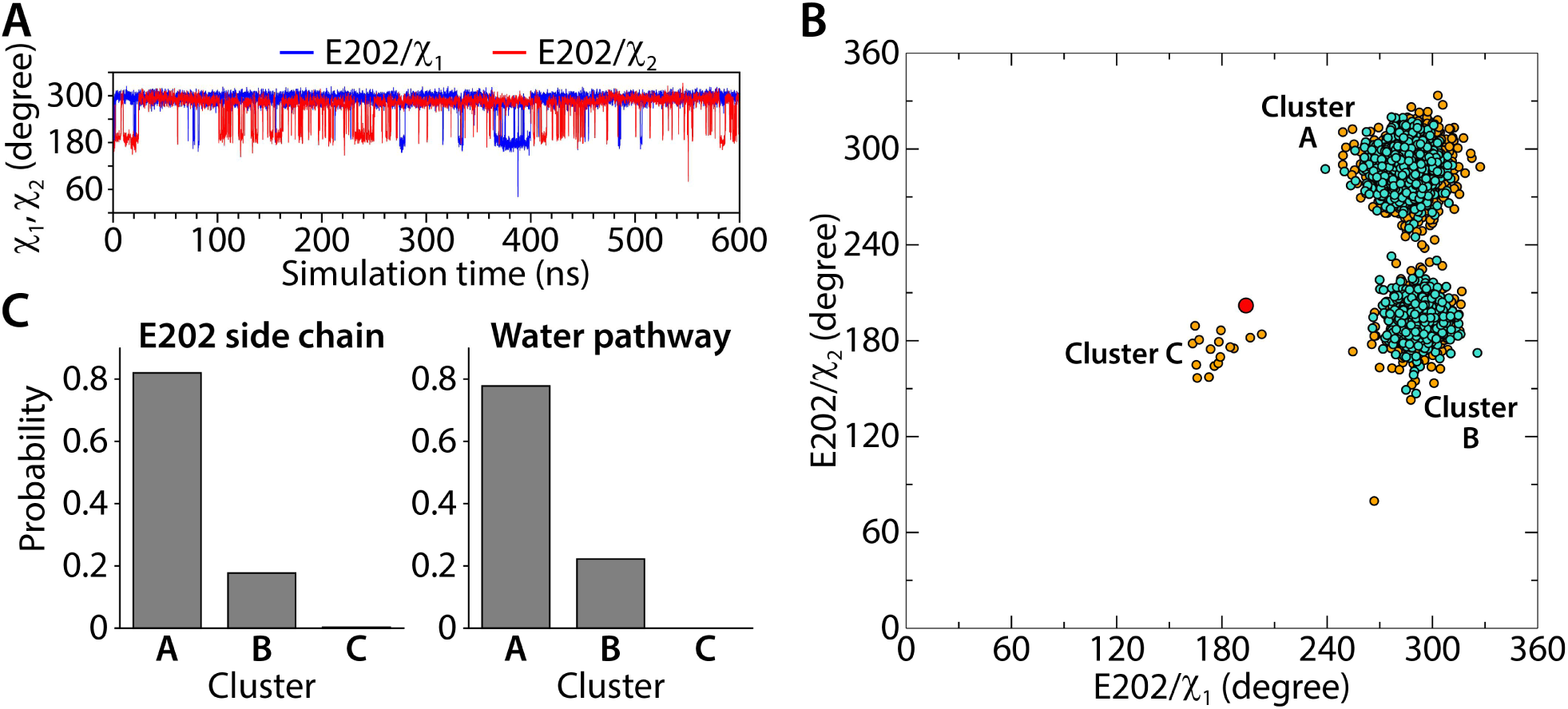
Conformational dynamics of the E202 side chain orientation (**A**) Traced dihedral angles of E202 during the simulation of the CLC-ec1 QQQ. (**B**) 2D scatter map of E202 dihedral angles reveals 3 clusters of side chain conformations sampled during the equilibrium simulation. Each dot represents one pair of the dihedral angles of E202 from snapshots taken at 100-ps intervals in the simulation. The frames with continuous water pathways are shown in cyan. The large red dot represents the starting position for E202, observed in the QQQ crystal structure. (**C**) The distribution of the side chain conformations of E202 over the trajectory (left panel) and water pathways (right panel, normalized to the total number of water pathways). Water pathways are only observed when E202 is in the cluster A or B conformation, rotated away from its starting position.

**Video 1** Dynamics of water pathways and the Gln_in_ sidechain in the QQQ mutant simulation. Representative simulation trajectory (250.5-337.5 ns) showing the dramatic hydration of the hydrophobic lumen by water penetration from the intracellular bulk. The Gln_ex_ (Q148) sidechain is accessible to the intracellular bulk through the continuous water pathways spontaneously and frequently formed during the simulation. The water pathway conformations undergo dynamical changes due to the sidechain orientations of Gln_in_. Protein helices are shown as orange ribbons. The Cl^−^ ion at the S_cen_ site is shown as a green sphere. Key amino acids in proximity to the water pathways or involved in Cl^−^-coordination are shown. See also Figure 5 and Figure 5 supplements.

## REFERENCES

Abraham, S.J., R.C. Cheng, T.A. Chew, C.M. Khantwal, C.W. Liu, S. Gong, R.K. Nakamoto, and M. Maduke. 2015. 13C NMR detects conformational change in the 100-kD membrane transporter ClC-ec1. J Biomol NMR. 61:209–226.

Accardi, A. 2015. Structure and gating of CLC channels and exchangers. J Physiol. 593:4129–4138.

Accardi, A., and C. Miller. 2004. Secondary active transport mediated by a prokaryotic homologue of ClC Cl- channels. Nature. 427:803–807.

Accardi, A., M. Walden, W. Nguitragool, H. Jayaram, C. Williams, and C. Miller. 2005. Separate ion pathways in a Cl-/H+ exchanger. J Gen Physiol. 126:563–570.

Basilio, D., K. Noack, A. Picollo, and A. Accardi. 2014. Conformational changes required for H(+)/Cl(-) exchange mediated by a CLC transporter. Nat Struct Mol Biol. 21:456–463.

Borbat, P.P., E.R. Georgieva, and J.H. Freed. 2013. Improved Sensitivity for Long-Distance Measurements in Biomolecules: Five-Pulse Double Electron-Electron Resonance. J Phys Chem Lett. 4:170–175.

Brandon, S., A.H. Beth, and E.J. Hustedt. 2012. The global analysis of DEER data. J Magn Reson. 218:93–104.

Caffrey, M., and V. Cherezov. 2009. Crystallizing membrane proteins using lipidic mesophases. Nat Protoc. 4:706–731.

Darden, T., D. York, and L. Pedersen. 1998. Particle mesh Ewald: An N·log(N) method for Ewald sums in large systems. J. Chem. Phys. 98:10089–10092.

Duster, A.W., C.M. Garza, B.O. Aydintug, M.B. Negussie, and H. Lin. 2019. Adaptive Partitioning QM/MM for Molecular Dynamics Simulations: 6. Proton Transport through a Biological Channel. J Chem Theory Comput. 15:892–905.

Dutzler, R., E.B. Campbell, M. Cadene, B.T. Chait, and R. MacKinnon. 2002. X-ray structure of a ClC chloride channel at 3.0 A reveals the molecular basis of anion selectivity. Nature. 415:287–294.

Dutzler, R., E.B. Campbell, and R. MacKinnon. 2003. Gating the selectivity filter in ClC chloride channels. Science. 300:108–112.

Elvington, S.M., C.W. Liu, and M.C. Maduke. 2009. Substrate-driven conformational changes in ClC-ec1 observed by fluorine NMR. EMBO J. 28:3090–3102.

Emsley, P., and K. Cowtan. 2004. Coot: model-building tools for molecular graphics. Acta Crystallogr D Biol Crystallogr. 60:2126–2132.

Engh, A.M., and M. Maduke. 2005. Cysteine accessibility in ClC-0 supports conservation of the ClC intracellular vestibule. J Gen Physiol. 125:601–617.

Estevez, R., B.C. Schroeder, A. Accardi, T.J. Jentsch, and M. Pusch. 2003. Conservation of chloride channel structure revealed by an inhibitor binding site in ClC-1. Neuron. 38:47–59.

Evans, P. 2006. Scaling and assessment of data quality. Acta Crystallogr D Biol Crystallogr. 62:72–82.

Faraldo-Gomez, J.D., and B. Roux. 2004. Electrostatics of ion stabilization in a ClC chloride channel homologue from Escherichia coli. J Mol Biol. 339:981–1000.

Feng, L., E.B. Campbell, Y. Hsiung, and R. MacKinnon. 2010. Structure of a eukaryotic CLC transporter defines an intermediate state in the transport cycle. Science. 330:635–641.

Feng, L., E.B. Campbell, and R. MacKinnon. 2012. Molecular mechanism of proton transport in CLC Cl-/H+ exchange transporters. Proc Natl Acad Sci U S A. 109:11699–11704.

Han, W., R.C. Cheng, M.C. Maduke, and E. Tajkhorshid. 2014. Water access points and hydration pathways in CLC H+/Cl- transporters. Proc Natl Acad Sci U S A. 111:1819–1824.

Hoopes, R.R., Jr., A.E. Shrimpton, S.J. Knohl, P. Hueber, B. Hoppe, J. Matyus, A. Simckes, V. Tasic, B. Toenshoff, S.F. Suchy, R.L. Nussbaum, and S.J. Scheinman. 2005. Dent Disease with mutations in OCRL1. Am J Hum Genet. 76:260–267.

Hoover, W.G. 1985. Canonical dynamics: Equilibrium phase-space distributions. Phys Rev A Gen Phys. 31:1695–1697.

Hu, H., S.A. Haas, J. Chelly, H. Van Esch, M. Raynaud, A.P. de Brouwer, S. Weinert, G. Froyen, S.G. Frints, F. Laumonnier, T. Zemojtel, M.I. Love, H. Richard, A.K. Emde, M. Bienek, C. Jensen, M. Hambrock, U. Fischer, C. Langnick, M. Feldkamp, W. Wissink-Lindhout, N. Lebrun, L. Castelnau, J. Rucci, R. Montjean, O. Dorseuil, P. Billuart, T. Stuhlmann, M. Shaw, M.A. Corbett, A. Gardner, S. Willis-Owen, C. Tan, K.L. Friend, S. Belet, K.E. van Roozendaal, M. Jimenez-Pocquet, M.P. Moizard, N. Ronce, R. Sun, S. O’Keeffe, R. Chenna, A. van Bommel, J. Goke, A. Hackett, M. Field, L. Christie, J. Boyle, E. Haan, J. Nelson, G. Turner, G. Baynam, G. Gillessen-Kaesbach, U. Muller, D. Steinberger, B. Budny, M. Badura-Stronka, A. Latos-Bielenska, L.B. Ousager, P. Wieacker, G. Rodriguez Criado, M.L. Bondeson, G. Anneren, A. Dufke, M. Cohen, L. Van Maldergem, C. Vincent-Delorme, B. Echenne, B. Simon-Bouy, T. Kleefstra, M. Willemsen, J.P. Fryns, K. Devriendt, R. Ullmann, M. Vingron, K. Wrogemann, T.F. Wienker, A. Tzschach, H. van Bokhoven, J. Gecz, T.J. Jentsch, W. Chen, H.H. Ropers, and V.M. Kalscheuer. 2016. X-exome sequencing of 405 unresolved families identifies seven novel intellectual disability genes. Mol Psychiatry. 21:133–148.

Huang, J., and A.D. MacKerell, Jr. 2013. CHARMM36 all-atom additive protein force field: validation based on comparison to NMR data. J Comput Chem. 34:2135–2145.

Humphrey, W., A. Dalke, and K. Schulten. 1996. VMD: visual molecular dynamics. J Mol Graph. 14:33-38, 27–38.

Hustedt, E.J., F. Marinelli, R.A. Stein, J.D. Faraldo-Gomez, and H.S. McHaourab. 2018. Confidence Analysis of DEER Data and Its Structural Interpretation with Ensemble-Biased Metadynamics. Biophys J. 115:1200–1216.

Iyer, R., C. Williams, and C. Miller. 2003. Arginine-agmatine antiporter in extreme acid resistance in Escherichia coli. J Bacteriol. 185:6556–6561.

Jentsch, T.J., and M. Pusch. 2018. CLC Chloride Channels and Transporters: Structure, Function, Physiology, and Disease. Physiol Rev. 98:1493–1590.

Jeschke, G. 2012. DEER distance measurements on proteins. Annu Rev Phys Chem. 63:419–446.

Jeschke, G., and Y. Polyhach. 2007. Distance measurements on spin-labelled biomacromolecules by pulsed electron paramagnetic resonance. Phys Chem Chem Phys. 9:1895–1910.

Jorgensen, W.L., J. Chandrasekhar, J.D. Madura, R.W. Impey, and M.L. Klein. 1983. Comparison of simple potential functions for simulating liquid water. J. Chem. Phys. 79:926–935.

Kabsch, W. 2010. Xds. Acta Crystallogr D Biol Crystallogr. 66:125–132.

Kasper, D., R. Planells-Cases, J.C. Fuhrmann, O. Scheel, O. Zeitz, K. Ruether, A. Schmitt, M. Poet, R. Steinfeld, M. Schweizer, U. Kornak, and T.J. Jentsch. 2005. Loss of the chloride channel ClC-7 leads to lysosomal storage disease and neurodegeneration. EMBO J. 24:1079–1091.

Khantwal, C.M., S.J. Abraham, W. Han, T. Jiang, T.S. Chavan, R.C. Cheng, S.M. Elvington, C.W. Liu, Mathews, II, R.A. Stein, H.S. McHaourab, E. Tajkhorshid, and M. Maduke. 2016. Revealing an outward-facing open conformational state in a CLC Cl(-)/H(+) exchange transporter. Elife. 5.

Klauda, J.B., R.M. Venable, J.A. Freites, J.W. O’Connor, D.J. Tobias, C. Mondragon-Ramirez, I. Vorobyov, A.D. MacKerell, Jr., and R.W. Pastor. 2010. Update of the CHARMM all-atom additive force field for lipids: validation on six lipid types. J Phys Chem B. 114:7830–7843.

Ko, Y.J., and W.H. Jo. 2010. Secondary water pore formation for proton transport in a ClC exchanger revealed by an atomistic molecular-dynamics simulation. Biophys J. 98:2163–2169.

Last, N.B., R.B. Stockbridge, A.E. Wilson, T. Shane, L. Kolmakova-Partensky, A. Koide, S. Koide, and C. Miller. 2018. A CLC-type F(-)/H(+) antiporter in ion-swapped conformations. Nat Struct Mol Biol. 25:601–606.

Lim, H.H., and C. Miller. 2009. Intracellular proton-transfer mutants in a CLC Cl-/H+ exchanger. J Gen Physiol. 133:131–138.

Lim, H.H., T. Shane, and C. Miller. 2012. Intracellular proton access in a Cl(-)/H(+) antiporter. PLoS Biol. 10:e1001441.

Lin, C.W., and T.Y. Chen. 2003. Probing the pore of ClC-0 by substituted cysteine accessibility method using methane thiosulfonate reagents. J Gen Physiol. 122:147–159.

Lloyd, S.E., S.H. Pearce, S.E. Fisher, K. Steinmeyer, B. Schwappach, S.J. Scheinman, B. Harding, A. Bolino, M. Devoto, P. Goodyer, S.P. Rigden, O. Wrong, T.J. Jentsch, I.W. Craig, and R.V. Thakker. 1996. A common molecular basis for three inherited kidney stone diseases. Nature. 379:445–449.

Lobet, S., and R. Dutzler. 2006. Ion-binding properties of the ClC chloride selectivity filter. EMBO J. 25:24–33.

Ludewig, U., M. Pusch, and T.J. Jentsch. 1996. Two physically distinct pores in the dimeric ClC-0 chloride channel. Nature. 383:340–343.

Matsumoto, M. 2007. Relevance of hydrogen bond definitions in liquid water. J Chem Phys. 126:054503.

Matulef, K., and M. Maduke. 2005. Side-dependent inhibition of a prokaryotic ClC by DIDS. Biophys J. 89:1721–1730.

Matulef, K., and M. Maduke. 2007. The CLC ‘chloride channel’ family: revelations from prokaryotes. Mol Membr Biol. 24:342–350.

Mayes, H.B., S. Lee, A.D. White, G.A. Voth, and J.M.J. Swanson. 2018. Multiscale Kinetic Modeling Reveals an Ensemble of Cl(-)/H(+) Exchange Pathways in ClC-ec1 Antiporter. J Am Chem Soc. 140:1793–1804.

McCoy, A.J., R.W. Grosse-Kunstleve, P.D. Adams, M.D. Winn, L.C. Storoni, and R.J. Read. 2007. Phaser crystallographic software. J Appl Crystallogr. 40:658–674.

Middleton, R.E., D.J. Pheasant, and C. Miller. 1996. Homodimeric architecture of a ClC-type chloride ion channel. Nature. 383:337–340.

Miller, C. 2006. ClC chloride channels viewed through a transporter lens. Nature. 440:484–489.

Miller, C., and W. Nguitragool. 2009. A provisional transport mechanism for a chloride channel-type Cl-/H+ exchanger. Philos Trans R Soc Lond B Biol Sci. 364:175–180.

Miller, C., and M.M. White. 1984. Dimeric structure of single chloride channels from Torpedo electroplax. Proc Natl Acad Sci U S A. 81:2772–2775.

Mishra, S., B. Verhalen, R.A. Stein, P.C. Wen, E. Tajkhorshid, and H.S. McHaourab. 2014. Conformational dynamics of the nucleotide binding domains and the power stroke of a heterodimeric ABC transporter. Elife. 3:e02740.

Morozenko, A., I.V. Leontyev, and A.A. Stuchebrukhov. 2014. Dipole Moment and Binding Energy of Water in Proteins from Crystallographic Analysis. J Chem Theory Comput. 10:4618–4623.

Murshudov, G.N., A.A. Vagin, and E.J. Dodson. 1997. Refinement of macromolecular structures by the maximum-likelihood method. Acta Crystallogr D Biol Crystallogr. 53:240–255.

Nguitragool, W., and C. Miller. 2007. CLC Cl /H+ transporters constrained by covalent cross-linking. Proc Natl Acad Sci U S A. 104:20659–20665.

Nishikawa, K.O., T.; Isogai, Y.; Saitô, N. 1972. Tertiary Structure of Proteins. I. Representation and Computation of the Conformations. Journal of the physical society of Japan. 32.

Nose, S. 1984. A unified formulation of the constant temperature molecular dynamics methods. J. Chem. Phys. 81:511–510.

Olsson, M.H., C.R. Sondergaard, M. Rostkowski, and J.H. Jensen. 2011. PROPKA3: Consistent Treatment of Internal and Surface Residues in Empirical pKa Predictions. J Chem Theory Comput. 7:525–537.

Park, E., and R. MacKinnon. 2018. Structure of the CLC-1 chloride channel from Homo sapiens. Elife. 7.

Park, K., B.C. Lee, and H.H. Lim. 2019. Mutation of external glutamate residue reveals a new intermediate transport state and anion binding site in a CLC Cl(-)/H(+) antiporter. Proc Natl Acad Sci U S A. 116:17345–17354.

Phillips, J.C., R. Braun, W. Wang, J. Gumbart, E. Tajkhorshid, E. Villa, C. Chipot, R.D. Skeel, L. Kale, and K. Schulten. 2005. Scalable molecular dynamics with NAMD. J Comput Chem. 26:1781–1802.

Phillips, S., A.E. Brammer, L. Rodriguez, H.H. Lim, A. Stary-Weinzinger, and K. Matulef. 2012. Surprises from an unusual CLC homolog. Biophys J. 103:L44–46.

Picollo, A., M. Malvezzi, J.C. Houtman, and A. Accardi. 2009. Basis of substrate binding and conservation of selectivity in the CLC family of channels and transporters. Nat Struct Mol Biol. 16:1294–1301.

Picollo, A., Y. Xu, N. Johner, S. Berneche, and A. Accardi. 2012. Synergistic substrate binding determines the stoichiometry of transport of a prokaryotic H(+)/Cl(-) exchanger. Nat Struct Mol Biol. 19:525–531, S521.

Robertson, J.L., L. Kolmakova-Partensky, and C. Miller. 2010. Design, function and structure of a monomeric ClC transporter. Nature. 468:844–847.

Rostkowski, M., M.H. Olsson, C.R. Sondergaard, and J.H. Jensen. 2011. Graphical analysis of pH-dependent properties of proteins predicted using PROPKA. BMC Struct Biol. 11:6.

Smart, O.S., J.G. Neduvelil, X. Wang, B.A. Wallace, and M.S. Sansom. 1996. HOLE: a program for the analysis of the pore dimensions of ion channel structural models. J Mol Graph. 14:354–360, 376.

Sobacchi, C., A. Villa, A. Schulz, and U. Kornak. 1993. CLCN7-Related Osteopetrosis. *In* GeneReviews((R)). M.P. Adam, H.H. Ardinger, R.A. Pagon, S.E. Wallace, L.J.H. Bean, K. Stephens and A. Amemiya, editors, Seattle (WA).

Stein, R.A., A.H. Beth, and E.J. Hustedt. 2015. A Straightforward Approach to the Analysis of Double Electron-Electron Resonance Data. Methods Enzymol. 563:531–567.

Stobrawa, S.M., T. Breiderhoff, S. Takamori, D. Engel, M. Schweizer, A.A. Zdebik, M.R. Bosl, K. Ruether, H. Jahn, A. Draguhn, R. Jahn, and T.J. Jentsch. 2001. Disruption of ClC-3, a chloride channel expressed on synaptic vesicles, leads to a loss of the hippocampus. Neuron. 29:185–196.

Stockbridge, R.B., H.H. Lim, R. Otten, C. Williams, T. Shane, Z. Weinberg, and C. Miller. 2012. Fluoride resistance and transport by riboswitch-controlled CLC antiporters. Proc Natl Acad Sci U S A. 109:15289–15294.

Veeramah, K.R., L. Johnstone, T.M. Karafet, D. Wolf, R. Sprissler, J. Salogiannis, A. Barth-Maron, M.E. Greenberg, T. Stuhlmann, S. Weinert, T.J. Jentsch, M. Pazzi, L.L. Restifo, D. Talwar, R.P. Erickson, and M.F. Hammer. 2013. Exome sequencing reveals new causal mutations in children with epileptic encephalopathies. Epilepsia. 54:1270–1281.

Walden, M., A. Accardi, F. Wu, C. Xu, C. Williams, and C. Miller. 2007. Uncoupling and turnover in a Cl-/H+ exchange transporter. J Gen Physiol. 129:317–329.

Wang, D., and G.A. Voth. 2009. Proton transport pathway in the ClC Cl-/H+ antiporter. Biophys J. 97:121–131.

Wang, Z., J.M.J. Swanson, and G.A. Voth. 2018. Modulating the Chemical Transport Properties of a Transmembrane Antiporter via Alternative Anion Flux. J Am Chem Soc. 140:16535–16543.

Winn, M.D., C.C. Ballard, K.D. Cowtan, E.J. Dodson, P. Emsley, P.R. Evans, R.M. Keegan, E.B. Krissinel, A.G. Leslie, A. McCoy, S.J. McNicholas, G.N. Murshudov, N.S. Pannu, E.A. Potterton, H.R. Powell, R.J. Read, A. Vagin, and K.S. Wilson. 2011. Overview of the CCP4 suite and current developments. Acta Crystallogr D Biol Crystallogr. 67:235–242.

Wu, E.L., X. Cheng, S. Jo, H. Rui, K.C. Song, E.M. Davila-Contreras, Y. Qi, J. Lee, V. Monje-Galvan, R.M. Venable, J.B. Klauda, and W. Im. 2014. CHARMM-GUI Membrane Builder toward realistic biological membrane simulations. J Comput Chem. 35:1997–2004.

Zhang, L., and J. Hermans. 1996. Hydrophilicity of cavities in proteins. Proteins. 24:433–438.

